# Evaluating metabolic and genomic data for predicting grain traits under high night temperature stress in rice

**DOI:** 10.1101/2022.10.27.514071

**Authors:** Ye Bi, Rafael Massahiro Yassue, Puneet Paul, Balpreet Kaur Dhatt, Jaspreet Sandhu, Thi Phuc Do, Harkamal Walia, Toshihiro Obata, Gota Morota

## Abstract

The asymmetric increase in average nighttime temperatures relative to increase in average daytime temperatures due to climate change is decreasing grain yield and quality in rice. Therefore, a better understanding of the impact of higher night temperature on single grain at whole genome level is essential for future development of more resilient rice. We investigated the utility of metabolites obtained from grains to classify high night temperature conditions of genotypes, and metabolites and single nucleotide polymorphisms to predict grain length, width, and perimeter phenotypes using a rice diversity panel. We found that the metabolic profiles of rice genotypes alone could be used to classify control and high night temperature conditions with high accuracy using random forest or extreme gradient boosting. The best linear unbiased prediction and BayesC showed greater metabolic prediction performance than machine learning models for grain-size phenotypes. Metabolic prediction was most effective for grain width, resulting in the highest prediction performance. Genomic prediction performed better than metabolic prediction. Integrating metabolites and genomics simultaneously in a prediction model slightly improved prediction performance. We did not observe a difference in prediction between the control and high night temperature conditions. Several metabolites were identified as auxiliary phenotypes that could be used to enhance the multi-trait genomic prediction of grain-size phenotypes. Our results showed that, in addition to single nucleotide polymorphisms, metabolites collected from grains offer rich information to perform predictive analyses, including classification modeling of high night temperature responses and regression modeling of grain size-related phenotypes in rice.

## Background

Sustainable increase in food production is paramount to meet the demands of the growing population. However, rising temperatures threaten the productivity of major food crops including rice (*O. Sativa*) (Peng et al., 2004; Wheeler and Von Braun, 2013; Zhao et al., 2017). Rice is the staple food in many countries, however, its productivity is threatened by an increase in the average minimum (nighttime) temperatures. There has been a greater rise in the rate of nighttime temperatures than that of daytime temperatures (Vose et al., 2005; Donat and Alexander, 2012; Xia et al., 2014). Recent studies have indicated that high night temperatures (HNT) negatively impact photosynthesis and respiration, and hence, rice grain yield (Welch et al., 2010; Peng et al., 2013; Jagadish et al., 2015; Wang et al., 2017; Impa et al., 2021). Importantly, HNT not only impacts grain yield-related traits but also grain width (Dhatt et al., 2021) and grain quality (Sreenivasulu et al., 2015; Wada et al., 2019) in rice. Given the increasing trend of global warming, understanding the variety of omic responses and their associations during grain development in rice is essential for improving its resilience to HNT.

Genomic prediction has been widely used to predict responses of plants and animals (Meuwissen et al., 2001). It is a powerful quantitative genetic approach to predict the genetic value of unphenotyped lines for diverse arrays of traits in rice (Bartholomé et al., 2022). In addition to DNA polymorphisms, metabolites have emerged as omics data sources that can be used to investigate biological responses. Plant metabolites play a multitude of critical roles in growth and development, and abiotic and biotic stress responses. The metabolites of plants are associated with nutrition, fragrance, and agronomic performance (Obata and Fernie, 2012). Differential metabolic abundance has been reported in rice grains between the control and HNT treatment (Dhatt et al., 2019), suggesting that the differences in biochemical or physiological signals between the two conditions are reflected in the metabolic profiles. Hence, it is worthwhile to investigate whether the metabolic profiles of genotypes alone can be used to classify control and HNT conditions. When single nucleotide polymorphism (SNP) data are used as predictors, a classification accuracy of 0.5 is expected because genomics is irrelevant to the presence or absence of HNT stress.

Prediction of phenotypes using metabolites, known as metabolic prediction, has been carried out in maize (Riedelsheimer et al., 2012; Guo et al., 2016; Westhues et al., 2017; Schrag et al., 2018) and rice (Xu et al., 2016), obtaining an encouraging result for its predictive ability. Metabolic prediction captures the molecular composition of a plant, such as changes in biochemical or physiological signals that influence phenotypes, which may not be directly explained by genomic prediction (Riedelsheimer et al., 2012). Thus, metabolic data can be used to evaluate plant growth- or stress-related phenotypes in response to HNT.

Despite its potential, metabolic responses to HNT stress and the use of metabolites as covariates for complex trait prediction have not been fully explored relative to genetic analysis in rice yet. Overall, we hypothesized that the inclusion of all available metabolites would be useful for metabolic classification and prediction in HNT studies. Therefore, the objectives of this study were threefold: 1) evaluate the classification ability of metabolites to distinguish HNT conditions, 2) compare the predictive ability of metabolic prediction, genomic prediction, and their multi-omic integration for grain-size phenotypes, and 3) investigate whether the use of metabolites as auxiliary phenotypes improves the predictive performance of multi-trait genomic prediction of grain-size phenotypes under control and HNT conditions.

## Materials and Methods

### Plant materials and growth conditions

Rice diversity panel 1 lines (Zhao et al., 2011) were phenotyped for grain length (major axis), grain width (minor axis), and grain perimeter in this study. Six seedlings per accession were transplanted into 4-inch pots containing natural soil. The HNT experiment was performed as previously described (Dhatt et al., 2021). Briefly, all the plants were grown under controlled conditions until flowering. When approximately 50% of the primary panicle completed fertilization, half of the plants from each accession were transferred to HNT conditions until maturity. All the plants were harvested at physiological maturity. Dehulled mature grains from primary panicles were scanned using an Epson Expression 12000 XL scanner (Epson America Inc., Los Alamitos, CA, USA) at a resolution of 600 dpi. Morphometric measurements, including grain length, width, and perimeter, were obtained from mature grains using the MATLAB software (Zhu et al., 2021). Morphometric phenotypes were adjusted for downstream genetic analyses by deriving the best linear unbiased estimators for each accession in each condition while accounting for replication. All the lines were genotyped using a high-density rice array (HDRA) of 700k SNP markers (McCouch et al., 2016). A total of 385,118 SNP markers were used for analysis after removing SNP markers with minor allele frequencies less than 0.05.

### Metabolic profiling

Five dehusked mature grains of each genotype were taken from the pool of all plant individuals and used for metabolite profiling. The grains were frozen and ground to fine powder by a ball mill (TissuelyzerII, Qiagen, Düsseldorf, Germany) at liquid nitrogen temperature. Around 50 mg of aliquot was weighed and used for the metabolite extraction and profiling using a 7200 GC-QTOF system (Agilent, Santa Clara, CA, USA) according to the protocol previously described (Wase et al., 2022). The chromatography peaks were annotated to metabolites according to the retention time and mass spectral information in the Fiehn Metabolomics database (Agilent). The peak heights of representative ions for individual metabolites were normalized by that of internal standard, ribitol (m/z 319), and the fresh weights of materials to determine relative metabolite contents. The retention time and representative ion m/z of each peak and the relative metabolite contents are found in the Supplementary Files. Relative metabolite abundance was corrected for run and experimental batch effects by treating them as random, separately for the control and HNT conditions.

### Statistical analyses

A total of 192 and 188 rice lines with phenotypes, genotypes, and metabolites were used for the control and HNT conditions, respectively. These lines consisted of tropical japonica (25.11%), temperate japonica (22.37%), indica (18.72%), aus (17.35%), admixed japonica (9.13%), aromatic (3.20%), admixed indica (2.74%), and admixed (1.38%) (McCouch et al., 2016). The utility of metabolic profiles to classify control and HNT conditions was evaluated. This was followed by a comparison of the predictive abilities of genomic prediction, metabolic prediction, and their combination for grain length, width, and perimeter. Finally, potential auxiliary metabolites that can be used to increase the multi-trait genomic prediction of grain-size phenotypes were explored under control and HNT conditions.

### Classification of HNT stress status

The following classification models were used to classify HNT stress from the control conditions based on the metabolic profiles of 380 (192 + 188) plants. Our hypothesis was that there is sufficient differential metabolic abundance between the control and HNT conditions that can be used to classify HNT stress status.

#### Logistic regression

Logistic regression (LR), which is built on the logit link function, models the probability that the metabolic profile of each plant belongs to the control or HNT stress status.

#### Support vector machine

Support vector machines (SVM) coupled with a radial basis function kernel was used to find the nonlinear separation boundary. The idea behind the SVM is to maximize the margin around the separating hyperplane (control or HNT status) by solving quadratic programming.

#### Random forest

Random forest (RF) is an ensemble learner based on numerous decision tree classifiers constructed from subsamples of the data. Each tree in the RF predicts the category (control or HNT status) under which a new plant in the testing set belongs. The final category was assigned to a new plant according to the majority vote.

#### Extreme gradient boosting

Extreme gradient boosting (XGBoost) is an ensemble machine learning framework that uses gradient boosted decision trees. Relative to the gradient boosting machine, XGBoost is faster and delivers higher prediction performance. We implemented LR, SVM, RF, and XGBoost in the caret R package (Kuhn, 2015).

### Metabolic prediction of grain-size phenotypes

In addition to the regression versions of SVM (i.e., support vector regression or SVR), RF, and XGBoost, ordinary least squares (OLS), best linear unbiased prediction (BLUP), and BayesC were used for the metabolic prediction of grain-size phenotypes.

#### Ordinary least squares

Metabolic OLS (MOLS) was constructed using metabolic abundance as a predictor in the OLS framework.

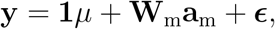

where **y** is a vector of phenotypes (grain length, grain width, and grain perimeter); **1** is the vector of ones; *μ* is the overall mean; **W**_m_ is a centered and standardized metabolic abundance matrix; **a**_m_ is a vector of fixed metabolic effect, and 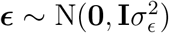, is a vector of residuals. Here 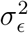 is the residual variance, and **I** is an identity matrix. The MOLS model was fitted using the lm function in R (R Core Team, 2022). This model was only used for metabolic prediction because the number of SNP markers was greater than the number of accessions in genomic prediction.

#### BayesC

A Bayesian shrinkage and variable selection model, BayesC (Kizilkaya et al., 2010), was used to estimate the metabolic effect using the following model.

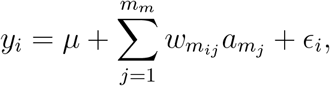

where *y_i_* is the vector of phenotypes for the *i*th accession; *m*_m_ is the total number of metabolites; *w_m_ij__* is the *j*th metabolic abundance of ith accession; and *a_m_j__* is the *j*th metabolic abundance effect. The prior of *a_m_j__* was assumed to folllow a mixture distribution

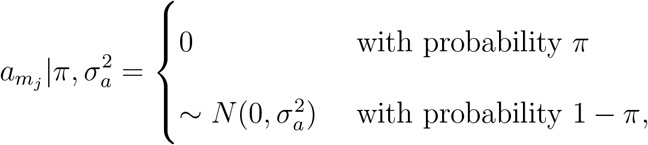

where 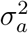 is the common metabolic abundance variance and *π* is a mixture proportion set to 0.99.

#### Metabolic best linear unbiased prediction

Best linear unbiased prediction regresses the vector of phenotypes on a kernel relationship matrix derived from the biological profiles of individuals (Morota and Gianola, 2014). The model considered for the metabolic best linear unbiased prediction (MBLUP) was

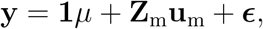

where **Z**_m_ is the incidence matrix relating metabolites to phenotypic records, **u**_m_ is the vector of the random metabolic values of the accessions, and **ϵ** is the vector of the residuals. The distributions of random metabolic effect was assumed to follow 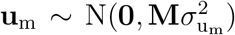, where **M** is the metabolic relationship matrix and 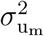 is the metabolic variance. The metabolic relationship matrix represents the similarity of metabolic profiles among accessions, which was computed as a function of the metabolic abundance cross-product 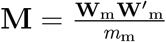.

### Genomic prediction of grain-size phenotypes

Performance of the metabolic prediction was compared with that of genomic best linear unbiased prediction (GBLUP), which is the most commonly used genomic prediction model (VanRaden, 2008). Here, metabolic abundance covariates were replaced with SNP marker covariates. The GBLUP model used was

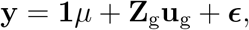

where **Z**_g_ is the incidence matrix relating gene content to phenotypic records and **u**_g_ is the vector of the random additive genetic values of the accessions. We assumed 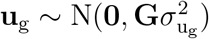, where 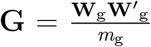 is the genomic relationship matrix; 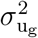 is the additive genetic variance; **W**_g_ is a centered and standardized gene content matrix; and *m*_g_ is the total number of SNP markers.

Metabolic 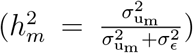 and genomic 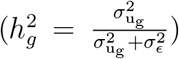 heritability estimates were ob-tained from MBLUP and GBLUP, respectively. These estimates can be interpreted as the proportion of phenotypic variance explained by metabolic or genomic relationship among lines.

Additionally, the exent of increased performance due to the integration of metabolites and SNP markers was evaluated by extending MBLUP and GBLUP via multiple kernel learning as follows (Baba et al., 2021).

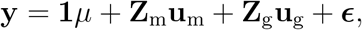

This approach was named integrated genomic metabolic best linear unbiased prediction (GMBLUP). Also, we performed the Mantel test to investigate whether the correlation between the **G** and **M** matrices is statistically different (Mantel, 1967).

The aforementioned BayesC, MBLUP, GBLUP, and GMBLUP were implemented in a Bayesian manner using the BGLR R package (Pérez and de los Campos, 2014). A flat prior was assigned to *μ*. The variance components, 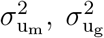, and 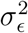 were drawn from a scaled inverse *χ*^2^ distribution with the degrees of freedom *v* = 5 and scale parameter *s* such that the prior means of variance components equal half of the phenotypic variance. A total of 30,000 Markov Chain Monte Carlo samples after 10,000 burn-in with a thinning rate of 10 were used to obtain the posterior means for all the unknowns.

### Multi-trait genomic prediction of grain-size phenotypes

We evaluated the gain in genomic prediction performance of grain-size phenotypes by fitting bivariate GBLUP, when metabolites were used as a correlated trait. We hypothesized that some metabolites could enhance the genomic prediction via a correlated response. All possible combinations of the phenotypes (target responses) and metabolites (auxiliary responses) were investigated. Genetic and residual variances in single-trait GBLUP were extended to the following variance-covariance structure.

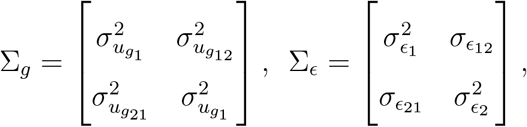

where subscripts 1 and 2 refer to the phenotype and metabolite, respectively. An inverse Wishart distribution was assigned to Σ_*g*_ and Σ_*ε*_ with degrees of freedom *v* = 4 and scale matrix *S* such that the prior means of Σ_*g*_ and Σ_*ε*_ equal half of the phenotypic variance. The bivariate GBLUP was fitted using 30,000 Markov chain Monte Carlo samples, 10,000 burn-ins, and a thinning rate of 10, implemented in the BGLR R package (Pérez-Rodríguez and de Los Campos, 2022).

### Cross-validation strategies

Repeated random subsampling cross-validation (CV) was used to evaluate the classification and prediction model performance. For classification, we first split the accessions into training (80%) and test (20%) sets separately for the control and HNT, so that each condition was represented in the training and testing sets equally (Figure 1A). The training set for each condition was further split into inner training and validation sets to fine-tune the hyperparameters. The inner training set was used for hyperparameter tuning using five-fold CV. The training sets from the control and HNT groups were combined to form a unified training set. The final model performance was evaluated in an independent testing set combined with the control and HNT conditions, which were never used in the model training. Repeated random sub-sampling CV for classification was performed 25 times. The accuracy of classification performance was derived as 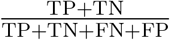, where TP, TN, FN, and FP are the number of accessions in the true positive, true negative, false negative, and false positive classes, respectively. Since the number of accessions in the control and HNT conditions were not exactly the same (192 and 188, respectively), we also evaluated classification performance using the F1 score and the area under a receiver operating characteristic (ROC) curve (AUC). The F1 score is robust to imbalanced data and is defined as the harmonic mean of the precision and recall 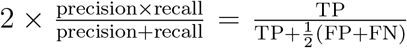. The AUC measures the area under the entire ROC curve, which plots the TP rate 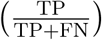 vs. the FP rate 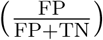. The accuracy and F1 score were derived using the Metrics R package (Hamner and Frasco, 2018) and the AUC was derived using the pROC R package (Robin et al., 2011). In addition to evaluating the utility of the whole metabolic profile for classification, we investigated the classification performance of random subsets of 10, 20, 30, 40, 50, and 60 metabolites by randomly reconstructing each subset 20 times.

**Figure 1:**
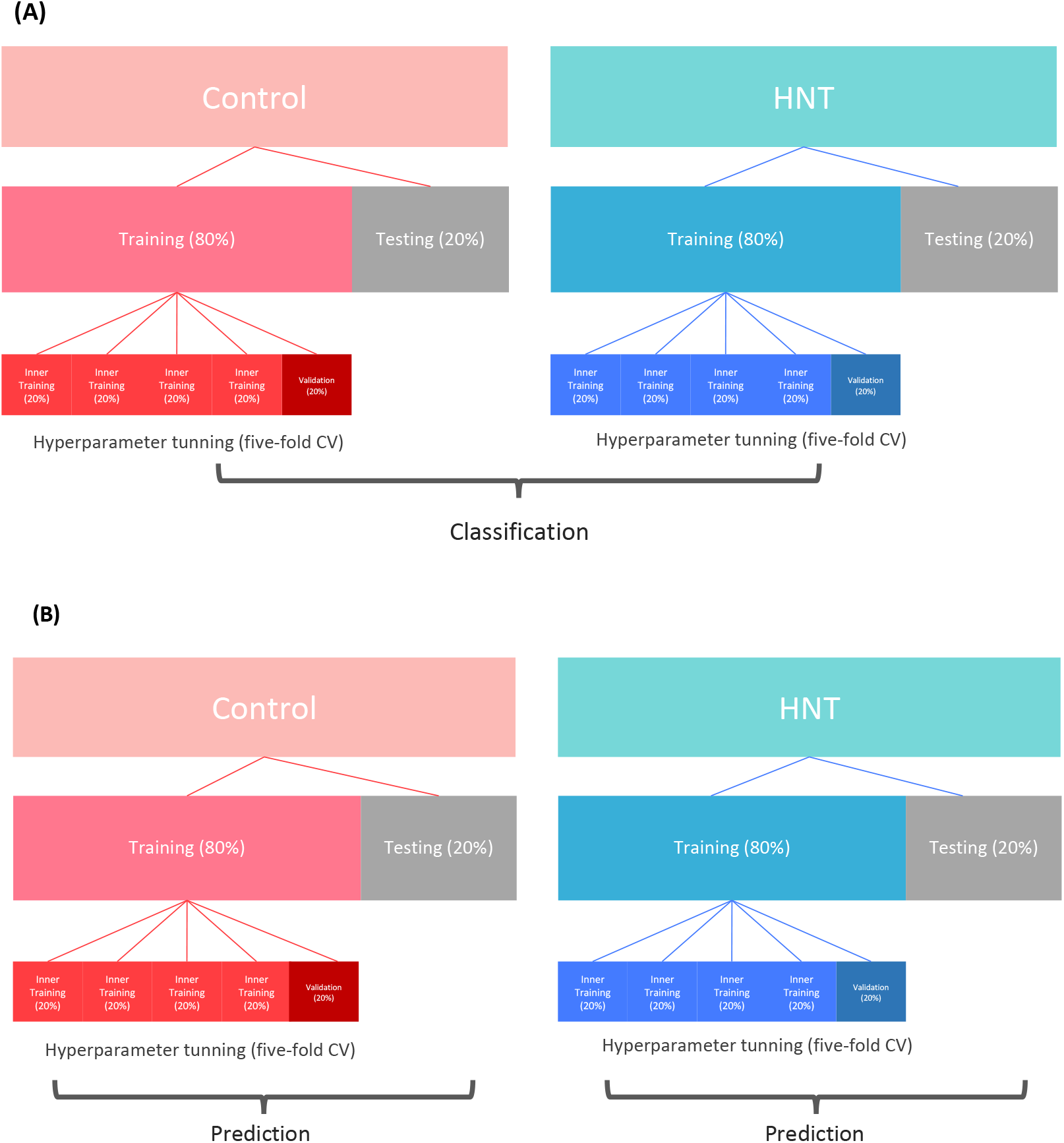
Cross-validation (CV) design for binary classification of high night temperature stress conditions (A) and metabolimic and genomic prediction of grain size related phenotypes (B).

The performance of metabolic and genomic predictions of grain-size phenotypes was evaluated similar to the classification, except that the predictions were performed separately for the control and HNT conditions (Figure 1B). The predictive performance of the models was assessed using Pearson correlation between the predictive values and phenotypes of the accessions. The repeated random subsampling CV for metabolic and genomic prediction was repeated 100 times. In metabolic prediction, we also evaluated whether metabolic effects estimated in one condition could be used to predict phenotypes in another. Specifically, we trained metabolites in the HNT and predicted phenotypes in the control and vice versa. This scenario investigate the transferability of the metabolic signal across stress conditions.

Two scenarios were considered for the multi-trait genomic prediction of grain-size phenotypes (Figure 2). Scenario 1 included splitting the accessions into training (80%) and testing (20%) sets. The models were trained in the training sets, and the predictive performance of the genomic prediction was evaluated in the remaining testing sets. Scenario 2 included the metabolic information of all accessions in a training set and assessed the genomic prediction performance of grain-size phenotypes using a testing set. The repeated random sub-sampling CV for multi-trait genomic prediction was repeated 25 times.

**Figure 2:**
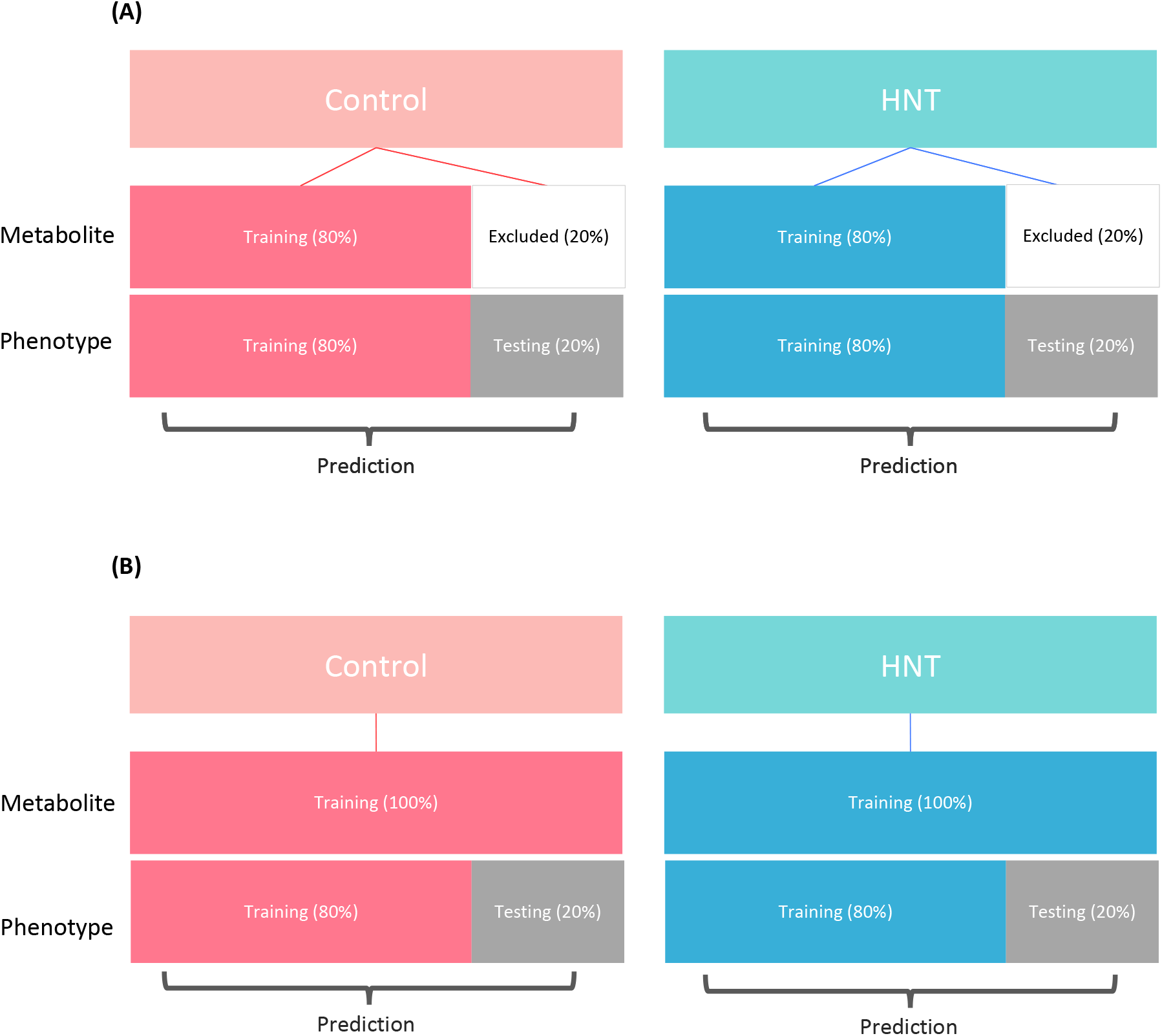
Cross-validation design for multi-trait prediction. Scenario 1 (A) and Scenario 2 (B).

### Data availability

Phenotypic and metabolic data used herein are available in the Supplementary Files at Figshare. Genotypic data regarding the rice accessions are available at the rice diversity panel website (http://www.ricediversity.org/). Scripts used in this work are publicly available in GitHub (https://github.com/yebigithub/VTUNL_Rice).

## Results

### Correlation analysis

Metabolic profiling of rice grains detected 73 metabolites (Table S1). Pairwise comparisons (*r*) of metabolic abundance revealed correlated metabolites (Figure S1). Under control conditions, four metabolites, citraconic acid, arabinose, lyxose, and ribose were positively associated with each other (|*r*| > 0.9) (Figure S1A). Pairs of leucine and valine, isoleucine and valine, isoleucine and leucine, and ornithine and citrulline also showed notable positive correlations. Under HNT, leucine was positively associated with isoleucine, arabinose, and ribose (|*r*| > 0.9). Citraconic acid was positively associated with ribose, adenine, and uridine (|*r*| > 0.9). Furthermore, pairs of dihydrouracil and asparagine, glutamine and asparagine, ribose and arabinose, and isomaltose and lyxose showed notable positive correlations (|*r*| > 0.9) (Figure S1A). When the metabolic abundance was expressed in terms of the ratio of control to HNT, arabinose and citraconic acid, arabinose and ribose appeared as two positively correlated metabolic pairs (|*r*| > 0.9) (Figure S2). Grain width was associated with many metabolites (Figure 3). In particular, proline, serine, aspartic acid, tyrosine, glucose, lysine, tryptophan, galactinol, and sucrose were positively correlated (*r* > 0.3), whereas trehalose was negatively correlated (*r* < −0.3) with grain width under control conditions (Figure 3A). In contrast, melibiose was positively correlated (*r* > 0.3), while trehalose was negatively correlated (*r* < −0.3) with grain width under HNT conditions (Figure 3B).

**Figure 3:**
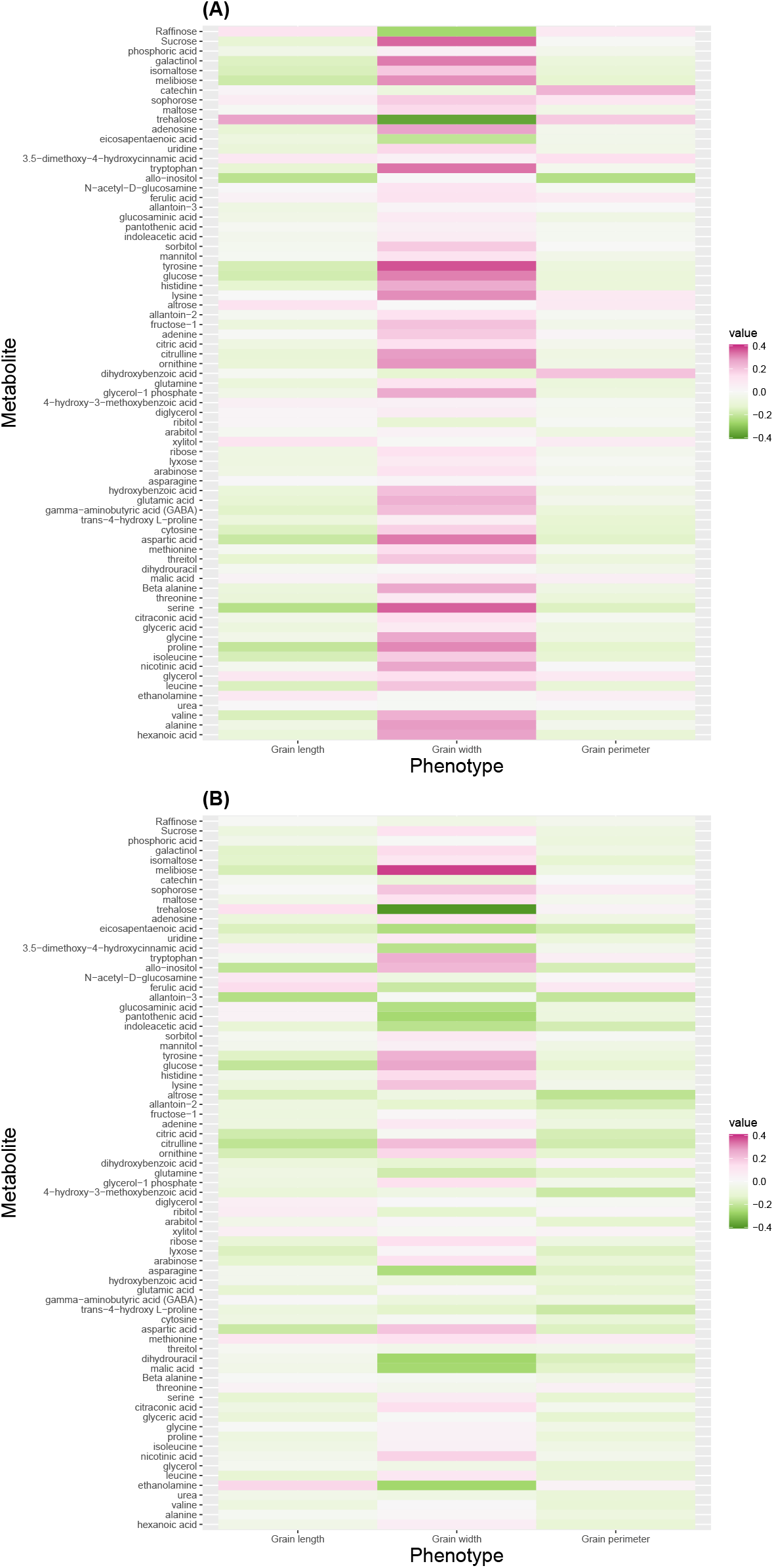
Correlation between metabolites and grain size phenotypes in control (A) and high night time temperature stress (B) conditions.

### Evaluation of metabolic classification performance

RF and XGBoost delivered the best classification accuracy equally, followed by SVM and LR (Figure 4). The means of classification for RF and XGBoost were both 0.98, suggesting that the metabolic profiles of the control and HNT conditions could be accurately used for classification. The mean SVM accuracy decreased moderately to 0.78. However, the LR classification performance was worse than that of a random classifier with a mean accuracy of 0.41 and a large CV uncertainty. Because slightly different number of accessions were used between the control and HNT conditions, the classification performance of the four models was evaluated using alternative measures. The F1 scores (Figure S3) and AUC (Figure S4) corroborated the accuracy results, suggesting that the classification accuracy performance obtained was robust. Classification accuracy was proportional to the number of metabolites included in the model (Figure S5). The opposite was observed in CV uncertainty, which was disproportional to the number of metabolites included in the model. A set of 10 metabolites alone achieved an accuracy above 0.8, albeit with a large CV uncertainty in RF and XGBoost. As the number of metabolites in the model increased, the accuracy increased and the CV uncertainty decreased. The accuracy of the 60 metabolites was slightly lower than that of all the metabolites. A set of 10 metabolites alone achieved an SVM accuracy of above 0.7, which gradually approached the accuracy achieved by the full set of metabolites. However, the LR did not follow this pattern. It consistently performed poorly regardless of the number of metabolites included in the model. The results of the F1 scores and AUC classification measures agreed with the classification accuracy (Figures S6 and S7).

**Figure 4:**
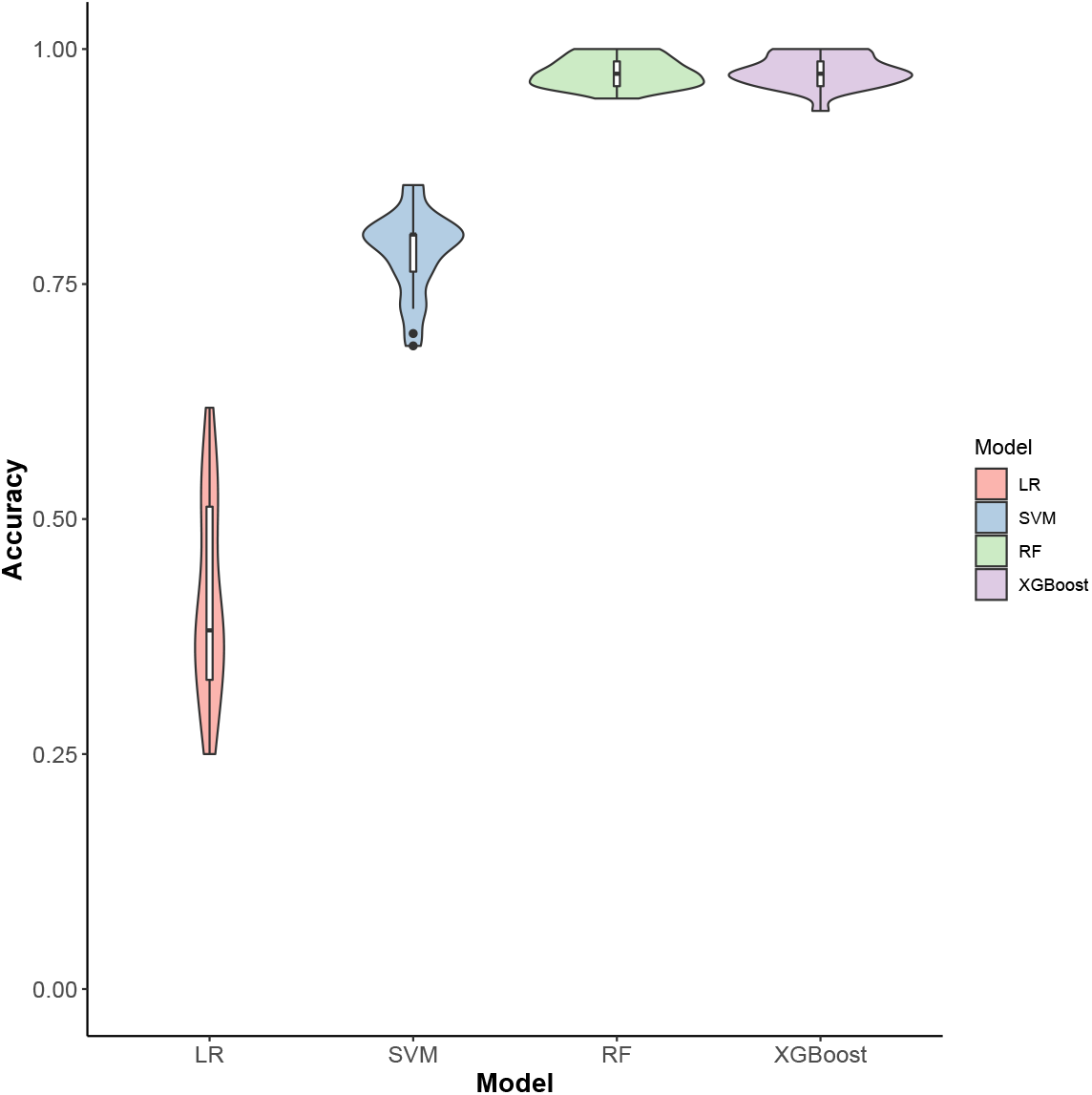
Classification accuracy of high night time temperature conditions (control and stress) using 73 metabolites. LR: logistic regression; SVM: support vector machine; RF: random forest; and XGBoost: extreme gradient boosting.

### Evaluation of metabolic prediction performance

The performance of the metabolic prediction of grain-size phenotypes using MOLS, RF, SVM, XGBoost, BayesC, and MBLUP is shown in Figures 5, 6, and 7. Points below the straight line indicate that the model shown on the x-axis performed better, whereas points above the straight line indicate the model shown on the y-axis performed better. BayesC and MBLUP were the equally best metabolic prediction models for grain length and delivered similar predicted values (Figure 5). Their mean predictive correlations were 0.35 and 0.33 in control and 0.33 and 0.31 in HNT conditions, respectively. However, although the means were similar, BayesC was better than MBLUP in 69% (control) and 80% (HNT) of the CV resampling runs. MOLS resulted in the worst performance, with mean predictive correlations of 0.20 (control) and 0.14 (HNT). The prediction performance of BayesC and MBLUP were higher than that of MOLS in more than 75% (control) and 84% (HNT) of the resampling runs. The metabolic prediction performance of the remaining models, RF, SVR, and XGBoost, was between that of BayesC or MBLUP and MOLS. For example, BayesC performed better than RF, SVR, and XGBoost in 84%, 77%, and 76% of the resampling runs in the control and 59%, 61%, and 59% of the resampling runs in the HNT.

**Figure 5:**
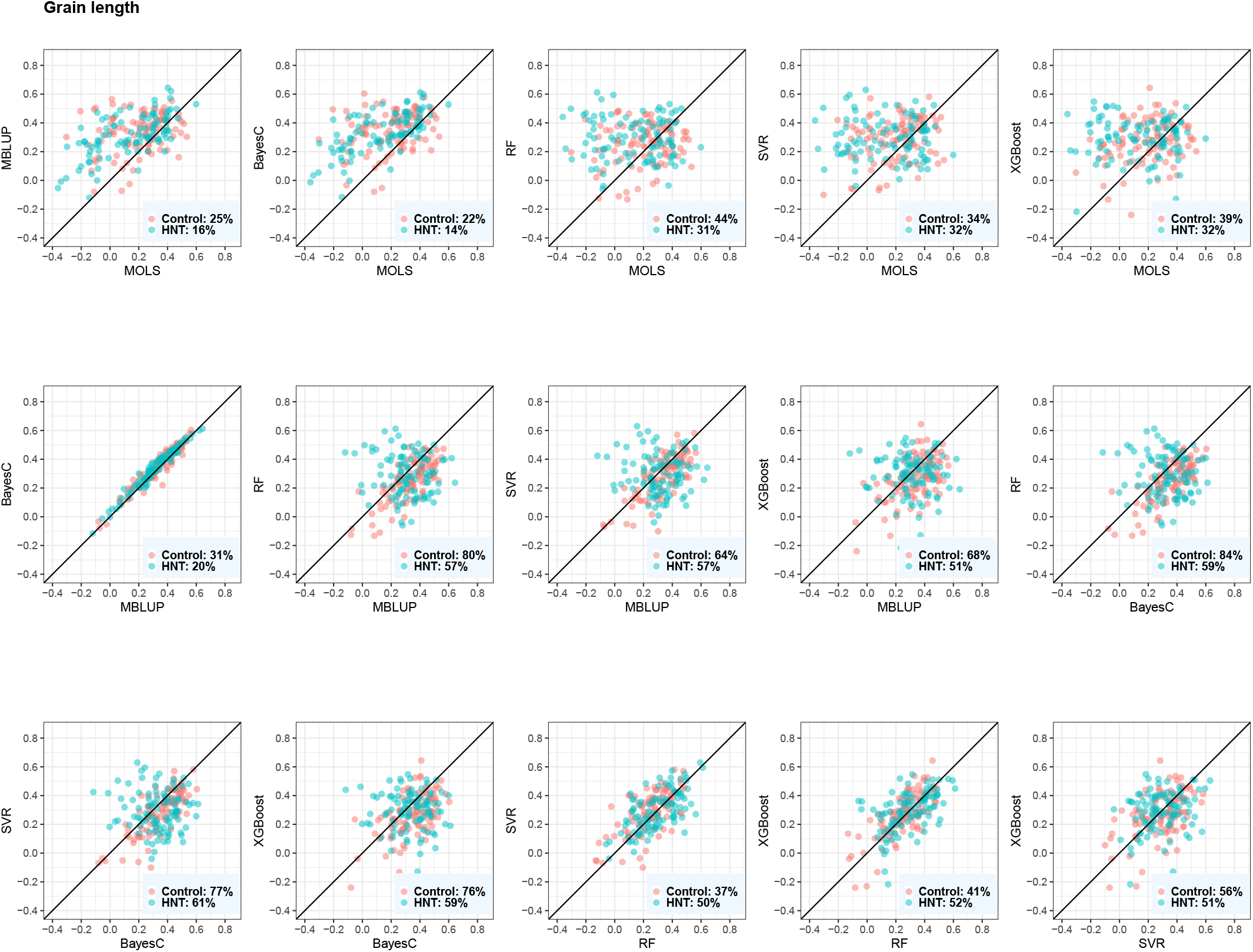
Predictive correlations of grain length using metabolic prediction in control and high night time temperature stress conditions. The percentages on the bottom right show the number of cross-validation resampling runs that the model on the x-axis performed better than the model on the y-axis. MOLS: metabolic ordinary least squares; MBLUP: metabolic best linear unbiased prediction; RF: random forests; SVR: support vector regression; and XGBoost: extreme gradient boosting.

For grain width measured under control conditions, RF was the best metabolic prediction model with a predictive correlation of 0.57, closely followed by MBLUP of 0.54 (Figure 6). RF was better than MOLS, BayesC, MBLUP, SVR, and XGBoost in 98%, 73%, 73%, 76%, and 73% of resampling runs, respectively. SVR, XGBoost, and BayesC performed equally well, and their predictive performance was better than that of MOLS. In the case of grain width measured under HNT conditions, BayesC and MBLUP equally delivered the best predictive correlation of 0.54, followed by SVR and XGBoost. For example, MBLUP showed a higher predictive performance than MOLS, BayesC, SVR, and XGBoost in 89%, 50%, 65%, and 70% of the resampling runs, respectively. Under both conditions, MOLS was the worst prediction machine.

**Figure 6:**
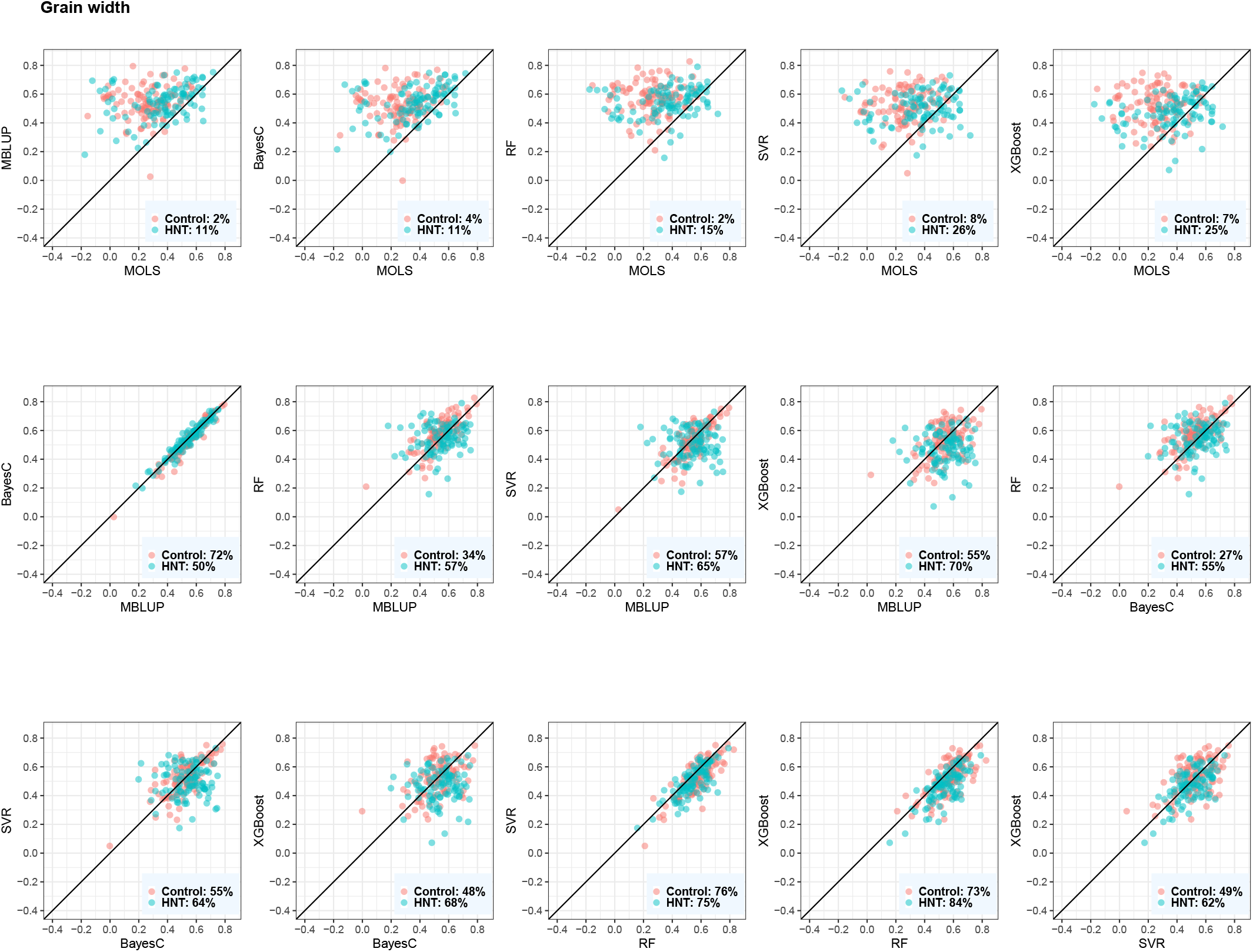
Predictive correlations of grain width using metabolic prediction in control and high night time temperature stress conditions. The percentages on the bottom right show the number of cross-validation resampling runs that the model on the x-axis performed better than the model on the y-axis. MOLS: metabolic ordinary least squares; MBLUP: metabolic best linear unbiased prediction; RF: random forests; SVR: support vector regression; and XGBoost: extreme gradient boosting.

For grain perimeter, BayesC consistently produced the best prediction (Figure 7). Its mean predictive correlations were 0.35 and 0.29 in the control and HNT conditions, respectively. BayesC performed better than MOLS, MBLUP, SVR, RF, and XGBoost in 88%, 75%, 69%, 89%, and 70% of the resampling runs in the control, whereas it performed better in 90%, 73%, 63%, 57%, and 51% of the resampling runs in the HNT.

**Figure 7:**
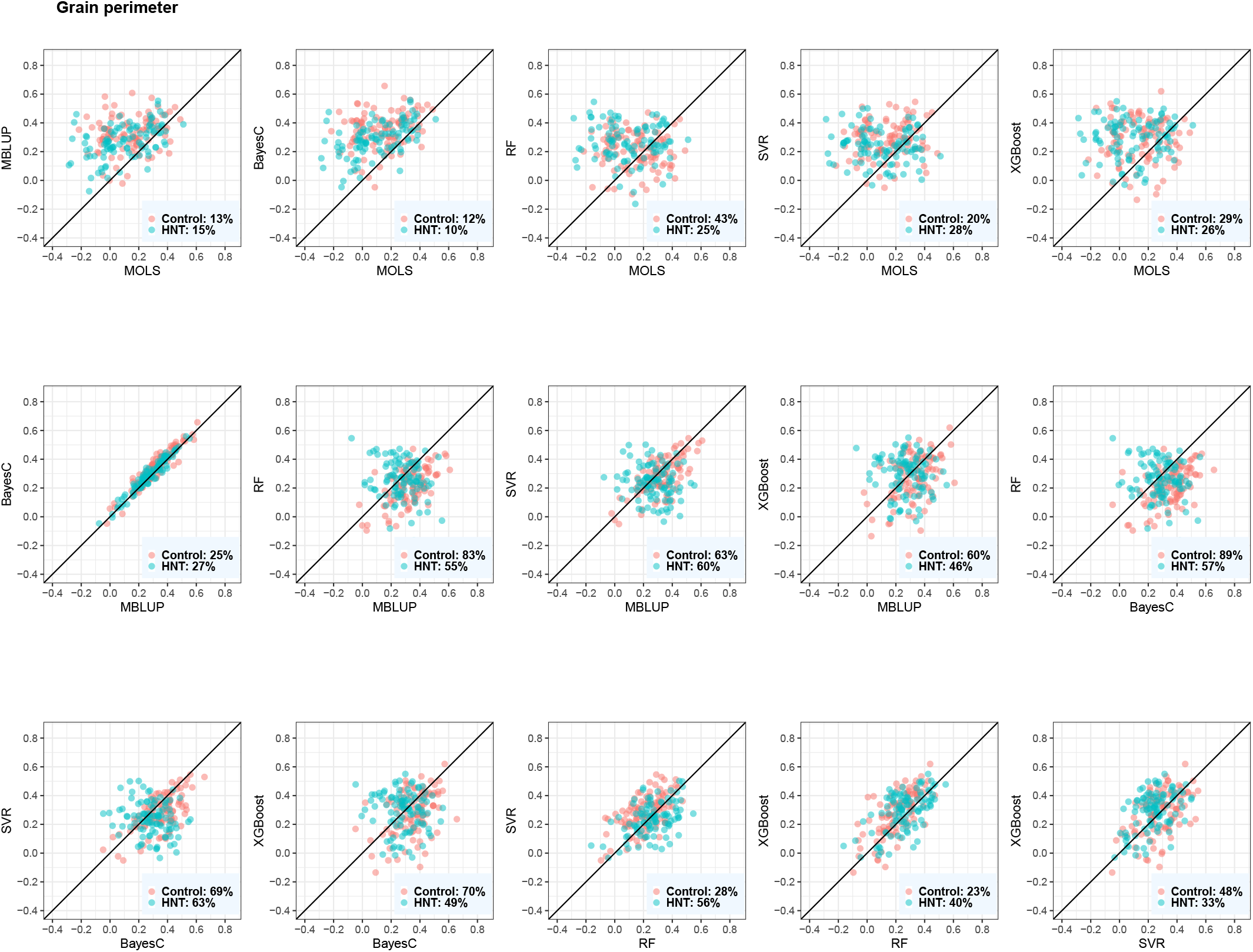
Predictive correlations of grain perimeter using metabolic prediction in control and high night time temperature stress conditions. The percentages on the bottom right show the number of cross-validation resampling runs that the model on the x-axis performed better than the model on the y-axis. MOLS: metabolic ordinary least squares; MBLUP: metabolic best linear unbiased prediction; RF: random forests; SVR: support vector regression; and XGBoost: extreme gradient boosting.

Overall, BayesC and MBLUP produced similar predicted metabolic values across the three phenotypes (Figures 5, 6, and 7). We obtained the highest prediction for grain width, whereas the predictive correlations of grain length and grain perimeter were similar and had slightly lower predictive outcomes. No significant difference was observed between the control and HNT conditions with respect to the predicted results. Additionally, we investigated whether the metabolic abundance obtained in one condition could be used to predict phenotypes under another condition. Overall, we found a decrease in metabolic prediction across HNT stress conditions (Figure S8). For example, when control phenotypes were predicted from HNT metabolites, we observed 23%, 4%, and 31% decrease in grain length, grain width, and grain perimeter, respectively, whereas when HNT phenotypes were predicted from control metabolites, we observed 18% and 4% decrease in grain width and grain perimeter, respectively. However, no decrease was observed in grain length.

### Evaluation of genomic prediction performance

The Mantel test showed that the correlation between **G** and **M** matrices are statistically different from each other. The performance of GBLUP and GMBLUP for grain-size phenotypes relative to that of MBLUP is shown in Figure 8. MBLUP was chosen to represent a metabolic prediction model because it performed well across the three traits under both conditions with a relatively faster computational time than BayesC. Overall, GBLUP consistently provided a better prediction than MBLUP in at least 98%, 87%, and 95% of CV resampling runs under both conditions for grain length, width, and perimeter, respectively. The mean predictive correlations were 0.64, 0.73, and 0.57 for grain length, width, and perimeter in control conditions, respectively, whereas 0.64, 0.67, 0.63 for grain length, width, and perimeter in HNT conditions, respectively. We observed mixed results for a genomics and metabolite integration model. In the case of grain length, GMBLUP did not improve the prediction when compared with that by GBLUP. GMBLUP performed better than GBLUP in only 48% (control) and 18% (HNT) of the resampling runs. However, in the case of grain width, GMBLUP performed better than GBLUP in 69% (control) and 72% (HNT) of the resampling runs. The results obtained for grain perimeter were mixed. Although the prediction performance of GMBLUP was better than that of GBLUP in 66% of the resampling runs under control conditions, GMBLUP performed better than GBLUP in only 37% of the resampling runs under HNT conditions. Overall, GMBLUP achieved a marginal gain in prediction than that achieved by GBLUP, with an average increase of 1.5%.

**Figure 8:**
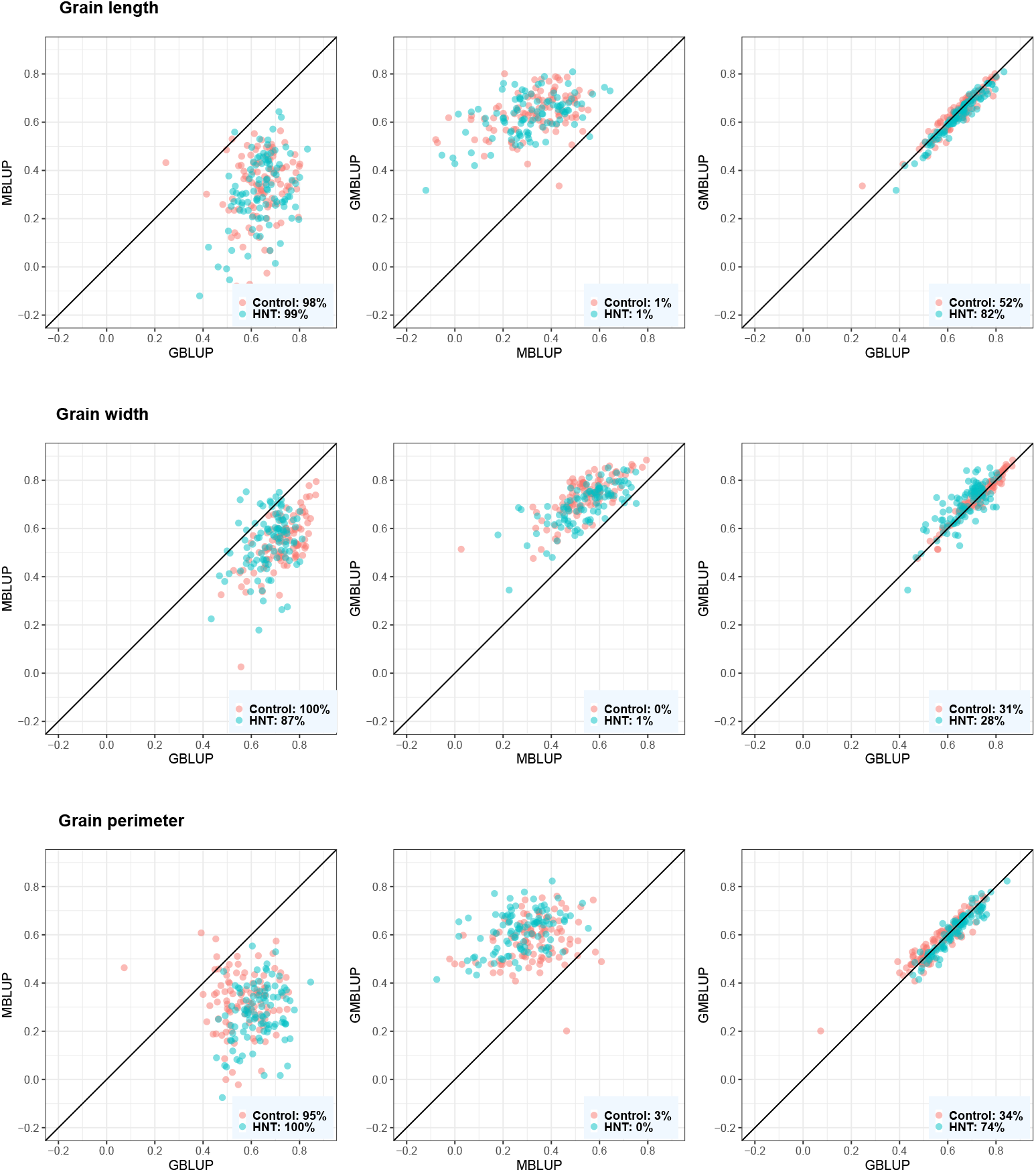
Predictive correlations of grain length, grain width, and grain perimeter using metabolic, genomic, and multi-omic prediction models in control and high night time temperature stress conditions. The percentages on the bottom right show the number of crossvalidation resampling runs that the model on the x-axis performed better than the model on the y-axis. MBLUP: metabolic best linear unbiased prediction; GBLUP: genomic best linear unbiased prediction; and GMBLUP: genomic and metabolic best linear unbiased prediction.

### Estimates of metabolic and genomic heritability

The metabolic and genomic heritabilities of grain-size phenotypes were estimated using MBLUP, GBLUP, and GMBLUP (Table 1). Under control conditions, genomic heritability estimates of grain length, width, and perimeter were similar, and explained at least 75% of the phenotypic variance. Grain width showed the highest metabolic heritability estimate, reaching more than half of the estimated genomic heritability. Grain length and grain perimeter showed lower metabolic heritability estimates than grain width. When the metabolites and SNP markers were fitted together, the majority of variations were captured by genomics. The estimates obtained from the HNT conditions were similar to those obtained from the control conditions. Grain width showed larger metabolic heritability estimates than grain length and perimeter. Genomics captured a large proportion of the variation when SNP markers and metabolites were simultaneously included in the model. However, the grain perimeter genomic heritability estimate was slightly lower than that of grain length and width.

**Table 1:** Metabolic heritability 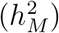 and genomic heritability 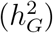 of grain size phenotypes in control and high night temperature stress conditions.

| Method | Grain length |  | Grain width |  | Grain perimeter |  |
| --- | --- | --- | --- | --- | --- | --- |
| | $h_G^2$ | $h_M^2$ | $h_G^2$ | $h_M^2$ | $h_G^2$ | $h_M^2$ |
| Control |  |  |  |  |  |  |
| GBLUP<sup>1</sup> | 0.78 | - | 0.76 | - | 0.75 | - |
| MBLUP<sup>2</sup> | - | 0.33 | - | 0.44 | - | 0.33 |
| GMBLUP<sup>3</sup> | 0.70 | 0.09 | 0.63 | 0.14 | 0.63 | 0.12 |
| High night temperature |  |  |  |  |  |  |
| GBLUP | 0.76 | - | 0.76 | - | 0.70 | - |
| MBLUP | - | 0.30 | - | 0.43 | - | 0.26 |
| GMBLUP | 0.67 | 0.08 | 0.60 | 0.15 | 0.61 | 0.08 |

<sup>1</sup> Genomic best linear unbiased prediction
<sup>2</sup> Metabolic best linear unbiased prediction
<sup>3</sup> Genomic metabolic best linear unbiased prediction

### Evaluation of multi-trait genomic prediction performance

The utility of metabolites as an auxiliary phenotype for multi-trait genomic prediction of grain-size phenotypes under control and HNT conditions was investigated in CV Scenarios 1 and 2. In Scenario 1, multi-trait GBLUP consistently produced a greater predictive correlation for grain width than single-trait GBLUP under the control conditions (Figure 9). All the metabolites included in the analyses contributed to increased prediction. We did not observe any increase in the prediction of grain length and perimeter. In Scenario 2, we identified at least one metabolite that increased the multi-trait genomic prediction performance for each trait (Figure 10). Three metabolites, glutamic acid, allantoin-2, and allantoin-3, increased multi-trait GBLUP prediction more than single-trait GBLUP for grain length under HNT conditions. No metabolites were found in grain length predictions under control conditions. A total of 46 metabolites improved predictions of grain width under control conditions. In particular, the gain in multi-trait GBLUP achieved by trehalose was statistically significant compared to that by single-trait GBLUP based on the paired one-sided t-test and paired one-sided Wilcoxon signed-rank test. Under HNT conditions, 11 metabolites, including ethanolamine, malic acid, dihydrouracil, asparagine, glutamine, allantoin-2, pantothenic acid, glucosaminic acid, ferulic acid, 3.5-dimethoxy-4-hydroxycinnamic acid, and trehalose, increased the genomic prediction performance for grain width. These metabolites were also identified in control conditions except for allantoin-2 and glucosaminic acid. Two metabolites, dihydroxybenzoic acid and catechin, increased the multi-trait GBLUP prediction for grain perimeter under control conditions. These values were statistically different from those of single-trait GBLUP. No metabolites were found in grain perimeter under HNT.

**Figure 9:**
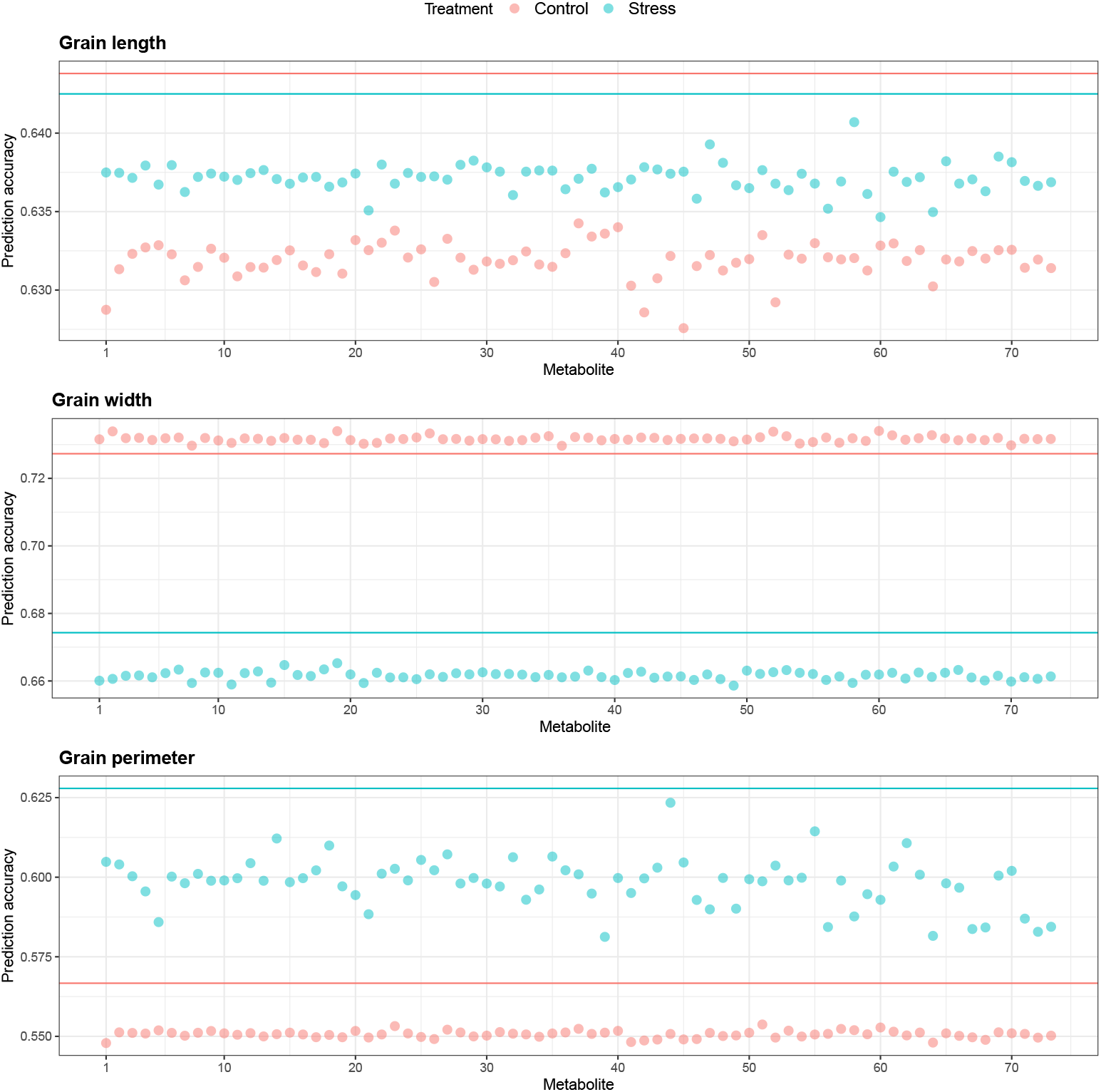
Predictive correlations of Scenario 1 multi-trait (bivariate) genomic prediction for grain length, grain width, and grain perimeter when metabolites were used as a secondary phenotype under control and high night time temperature stress conditions. The horizontal lines indicate the predictive correlations of single-trait genomic prediction.

**Figure 10:**
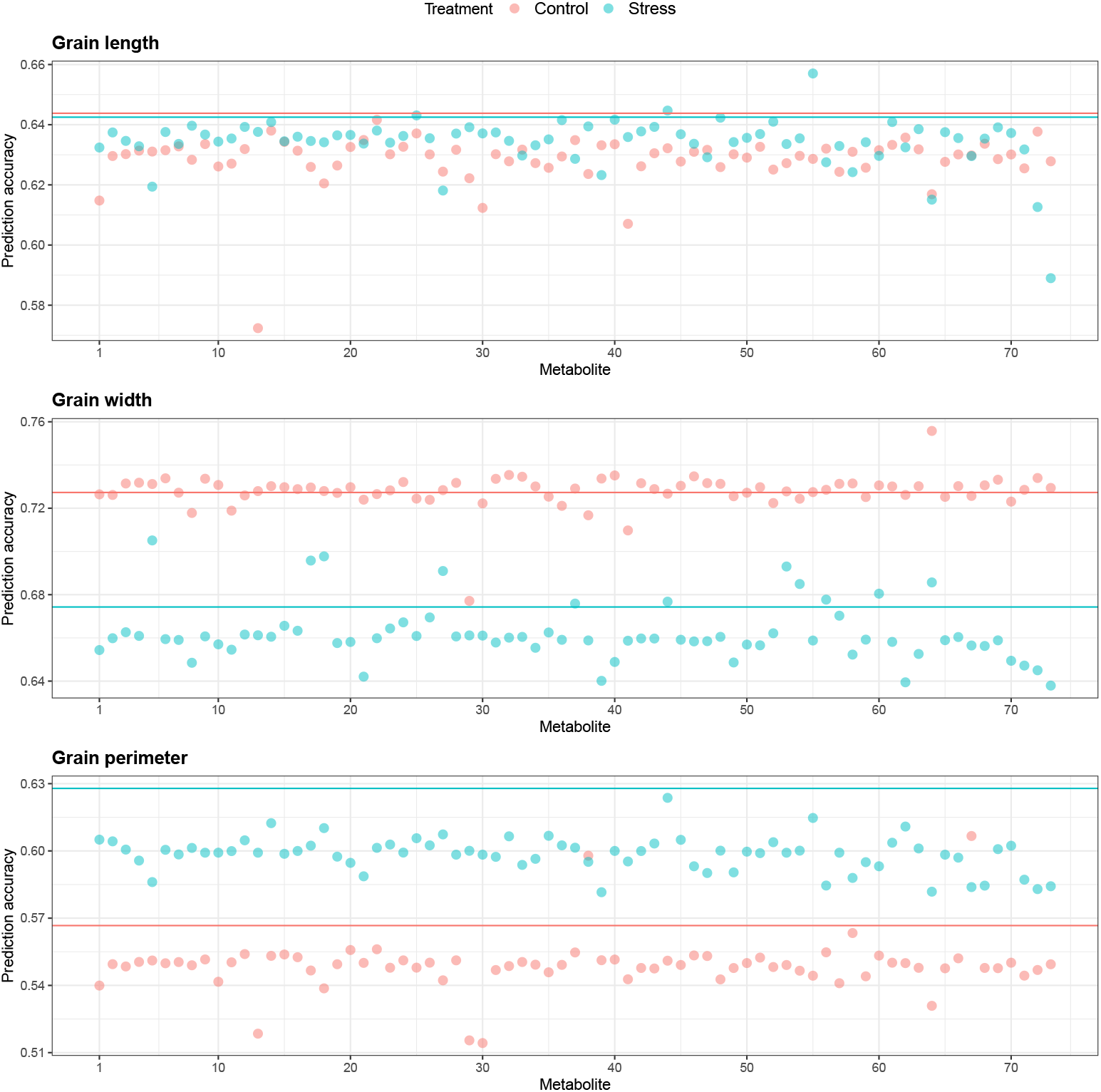
Predictive correlations of Scenario 2 multi-trait (bivariate) genomic prediction for grain length, grain width, and grain perimeter when metabolites were used as a secondary phenotype under control and high night time temperature stress conditions. The horizontal lines indicate the predictive correlations of single-trait genomic prediction.

## Discussion

Advances in genomics and metabolic profiling have provided a new resource for studying HNT responses in rice. In this study, we evaluated the utility of metabolites for classifying HNT stress conditions and predicting grain-size-related phenotypes. In regression modeling, the performance of metabolic prediction and usefulness of metabolites as auxiliary phenotypes were evaluated in the context of genomic prediction. We found that several pairs of metabolites were correlated under the control and HNT conditions. Among the three phenotypes investigated, metabolite abundance was strongly associated with grain width. Mostly amino acids and sugars were correlated with grain width under the control condition (Figure 3), indicating the relationship between grain shapes and carbohydrate and protein metabolism. The correlation between tryptophan and seed longevity has been also reported (Ren et al., 2020), supporting our result. Under the HNT condition, oligosaccharides showed correlations with grain width, which may indicate that carbohydrate metabolism plays a crucial role in determining grain shape. Interestingly, carbohydrates are a major class of metabolites which are affected by HNT in our previous studies of cereals grains (Dhatt et al., 2019; Impa et al., 2019). The correlation may reflect the changes in carbohydrate metabolism in rice grains under HNT. However, it should also be considered that differences in grain shapes among cultivars could affect metabolite composition due to the difference in the ratios of cell types with varying metabolite compositions. The relationships between metabolite accumulation and grain shapes must be carefully assessed in future experiments.

Grain length and width are prominent grain size factors that substantially impact the rice grain yield parameters (Olsen, 2004; Xing et al., 2010; Huang et al., 2013). Grain length is known to be directed by the elongation of the pericarp tissue, defined at the early stages of grain development (Lizana et al., 2010; Pielot et al., 2015). In contrast, the grain width is largely determined by cell division and proliferation of the endosperm tissue (after the fertilization event) and is a driving force for the sucrose allocation to be used for endosperm development and grain storage reserves production (Brocklehurst, 1977; Martinez-Carrasco and Thorne, 1979). The endosperm cell number and proliferation (determinant of grain width) are affected by the supply of photoassimilates (sucrose) during the active grain filling stage, thus, substantially influencing the final grain weight parameters (Brocklehurst, 1977). Likewise, another study reported that final grain weight is highly correlated with the grain width than the grain length in the diverse winter wheat population (Philipp et al., 2018). The improvement in grain width is stated to be one of the major causes leading to increment in the final grain weight parameters for the elite wheat varieties (Philipp et al., 2018), signifying the importance of this phenotypic trait for the enhancement of crop yields. Additionally, previous studies have reported that deviation from optimal temperature conditions alters the endosperm cellularization timing in rice, causing a detrimental impact on the final grain size parameters (Chen et al., 2016; Folsom et al., 2014). Therefore, it is likely that the prolonged occurrence of HNT during grain filling stages impairs the sucrose allocation in the endosperm cells, leading to a more negative impact on the grain width than grain length, which is established before grain width. Furthermore, the genetic determinants regulating these two grain size traits (grain length and width) have distinct responses to temperature abnormalities within rice diversity panel 1 accessions (Dhatt et al., 2021). Only a few accessions of rice diversity panel 1 retained both grain length and width under HNT, signifying unique genetic regulation for these grain size traits in rice (Dhatt et al., 2021).

### Utility of metabolites for classification

Using all available metabolites resulted in the appropriate classification of HNT conditions with high accuracy when suitable classification models, such as RF or XGBoost, were used. This suggests that there is a differential metabolic abundance between control and HNT conditions. Logistic regression, which is a simpler classification model, was not sufficient to distinguish the signals between the control and HNT conditions. A random subset of only 10 metabolites produced moderate classification accuracy. However, this accuracy was unstable with high uncertainty. Increasing the number of metabolites contributed to making the classification more robust than increasing its accuracy. The accuracy achieved from a random subset of 60 metabolites was similar to that achieved from a full set of metabolites. This implies that most metabolites are altered during HNT and contributes to increasing the classification power. Although there are no previous reports investigating the classification performance of metabolites to distinguish between control and HNT conditions, our results showed that we can obtain reasonable classification accuracy.

### Utility of metabolites for prediction and heritability analysis compared to SNP markers

Overall, BayesC and MBLUP showed relatively high and stable predictive correlations for grain length, width, and perimeter, suggesting that prediction models commonly used in genomic prediction are equally applicable to metabolic abundance data. In particular, BLUP appeared to be the most efficient method in terms of predictability and computational time. The extent of predictive correlations ranged from low to moderate. As expected from the correlation analysis, we observed the greatest predictive correlation for grain width. However, genomic prediction performance based on a 700k array was consistently better than that of metabolites. Our results agree with those of previous studies in oats, which found that metabolic prediction was not superior to genomic prediction for agronomic traits, including seed length and width (Hu et al., 2021). Similarly, no improvement was observed in 1000-grain weight and grain number per plant in hybrid rice (Xu et al., 2016; Wang et al., 2019; Xu et al., 2021). In contrast, metabolic prediction has been reported to be better than genomic prediction for oat fatty acids (Hu et al., 2021). In addition, other related studies reported lower or comparable predictive performance of metabolites relative to SNP markers for agronomic traits related to maturity, morphology, bioenergy, or yield-related in inbred maize (Guo et al., 2016; Xu et al., 2017), hybrid maize (Riedelsheimer et al., 2012; Westhues et al., 2017; Schrag et al., 2018), and hybrid wheat (Zhao et al., 2015). Simultaneous integration of metabolites and SNP markers in a single model moderately improved predictions for grain width in control and HNT conditions and grain perimeter in control conditions. This is partly in line with Hu et al. (2021), who reported a gain in GMBLUP prediction for seed length and width in oats. On an average, the estimates of metabolic heritability were approximately half of those of genomic heritability. A smaller number of available metabolites may lead to lower estimates. Another reason could be that the metabolite-phenotype relationship is affected by the time of measurement in metabolite profiling. Unlike SNP markers, metabolite abundance varies with plant developmental stages. Nevertheless, metabolites explained a greater proportion of variability when compared at the per metabolite or per SNP level. For example, in grain width, each metabolite explained 0.6% of the metabolic heritability, whereas each SNP explained 0.0001% of the genomic heritability.

### Utility of metabolites as auxiliary phenotypes in genomic prediction

We identified several metabolites that contributed to increased multi-trait genomic prediction of grain-size-related phenotypes. In particular, several metabolites aided the genomic prediction of grain width. Trehalose is a metabolite whose levels in grains were correlated with grain width (Figures 3, S1, and S2) and significantly contributed to the grain width prediction (Figures 9 and 10) in both control and HNT conditions. Trehalose is closely related to trehalose-6-phosphate (T6P), a signaling metabolite involved in the regulations of photosynthesis, carbon partitioning, and reproductive development (Oszvald et al., 2018; Ponnu et al., 2011; Smeekens, 2015). T6P accumulation correlates with the levels of sucrose and other major sugars in wheat spikes (Martínez-Barajas et al., 2011). Therefore, it is likely that the differences in trehalose and T6P metabolism that appeared as the accumulation of trehalose in grains among rice cultivars affected grain development and shape. It is also interesting to note that trehalose accumulation is affected by HNT in developing rice and wheat grains (Dhatt et al., 2019; Impa et al., 2019). The correlation between trehalose and grain width and the trehalose contribution to the prediction may suggest the influences of trehalose metabolism in determining grain shape under HNT conditions.

## Conclusions

This study showed that the metabolic profiles of rice genotypes could be used solely to classify control and HNT conditions with high accuracy. Although the metabolic prediction of grainsize phenotypes was low to moderate, they were the most effective for grain width. Genomic prediction delivered better results than metabolic prediction. The simultaneous integration of metabolites and genomics in a single statistical model yielded a minor improvement in prediction. No significant differences in metabolic or genomic predictions were observed between the control and HNT conditions. We identified several metabolites that can be used to enhance multi-trait genomic prediction as auxiliary phenotypes. These metabolites could be candidates for further studies to understand their role in the tolerance to HNT. Taken together, this study demonstrates the usefulness of rice metabolites for classification and prediction tasks under control and HNT conditions.

## Author contribution statement

PP, BKD, JS, and HW performed the high night temperature stress experiments. TPD and TO performed the metabolic analysis. YB analyzed the data. RMY supported YB on the data analysis. YB and GM drafted the manuscript. GM supervised the study and designed the data analysis. RMY, PP, BKD, JS, TPD, HW, and TO edited the manuscript.

## Acknowledgments

TPD acknowledges support from the Fulbright visiting scholar program.

## References

Baba, T., Pegolo, S., Mota, L. F., Peñagaricano, F., Bittante, G., Cecchinato, A., and Morota, G. (2021). Integrating genomic and infrared spectral data improves the prediction of milk protein composition in dairy cattle. Genetics Selection Evolution, 53(1):1–14.

Bartholomé, J., Prakash, P. T., and Cobb, J. N. (2022). Genomic prediction: progress and perspectives for rice improvement. Complex Trait Prediction, pages 569–617.

Brocklehurst, P. (1977). Factors controlling grain weight in wheat. Nature, 266(5600):348–349.

Chen, C., Begcy, K., Liu, K., Folsom, J. J., Wang, Z., Zhang, C., and Walia, H. (2016). Heat stress yields a unique mads box transcription factor in determining seed size and thermal sensitivity. Plant physiology, 171(1):606–622.

Dhatt, B. K., Abshire, N., Paul, P., Hasanthika, K., Sandhu, J., Zhang, Q., Obata, T., and Walia, H. (2019). Metabolic dynamics of developing rice seeds under high night-time temperature stress. Frontiers in Plant Science, 10:1443.

Dhatt, B. K., Paul, P., Sandhu, J., Hussain, W., Irvin, L., Zhu, F., Adviento-Borbe, M. A., Lorence, A., Staswick, P., Yu, H., et al. (2021). Allelic variation in rice fertilization independent endosperm 1 contributes to grain width under high night temperature stress. New Phytologist, 229(1):335–350.

Donat, M. G. and Alexander, L. V. (2012). The shifting probability distribution of global daytime and night-time temperatures. Geophysical Research Letters, 39(14).

Folsom, J. J., Begcy, K., Hao, X., Wang, D., and Walia, H. (2014). Rice fertilization-independent endosperm1 regulates seed size under heat stress by controlling early endosperm development. Plant Physiology, 165(1):238–248.

Guo, Z., Magwire, M. M., Basten, C. J., Xu, Z., and Wang, D. (2016). Evaluation of the utility of gene expression and metabolic information for genomic prediction in maize. Theoretical and applied genetics, 129(12):2413–2427.

Hamner, B. and Frasco, M. (2018). Metrics: Evaluation Metrics for Machine Learning. R package version 0.1.4.

Hu, H., Campbell, M. T., Yeats, T. H., Zheng, X., Runcie, D. E., Covarrubias-Pazaran, G., Broeckling, C., Yao, L., Caffe-Treml, M., Gutiérrez, L., et al. (2021). Multi-omics prediction of oat agronomic and seed nutritional traits across environments and in distantly related populations. Theoretical and Applied Genetics, 134(12):4043–4054.

Huang, R., Jiang, L., Zheng, J., Wang, T., Wang, H., Huang, Y., and Hong, Z. (2013). Genetic bases of rice grain shape: so many genes, so little known. Trends in plant science, 18(4):218–226.

Impa, S. M., Raju, B., Hein, N. T., Sandhu, J., Prasad, P. V., Walia, H., and Jagadish, S. K. (2021). High night temperature effects on wheat and rice: Current status and way forward. Plant, Cell & Environment, 44(7):2049–2065.

Impa, S. M., Sunoj, V. J., Krassovskaya, I., Bheemanahalli, R., Obata, T., and Jagadish, S. K. (2019). Carbon balance and source-sink metabolic changes in winter wheat exposed to high night-time temperature. Plant, Cell & Environment, 42(4):1233–1246.

Jagadish, S., Murty, M., and Quick, W. (2015). Rice responses to rising temperatures-challenges, perspectives and future directions. Plant, cell & environment, 38(9):1686–1698.

Kizilkaya, K., Fernando, R., and Garrick, D. (2010). Genomic prediction of simulated multibreed and purebred performance using observed fifty thousand single nucleotide polymorphism genotypes. Journal of animal science, 88(2):544–551.

Kuhn, M. (2015). Caret: classification and regression training. Astrophysics Source Code Library, pages ascl–1505.

Lizana, X. C., Riegel, R., Gomez, L. D., Herrera, J., Isla, A., McQueen-Mason, S. J., and Calderini, D. F. (2010). Expansins expression is associated with grain size dynamics in wheat (triticum aestivum l.). Journal of experimental botany, 61(4):1147–1157.

Mantel, N. (1967). The detection of disease clustering and a generalized regression approach. Cancer research, 27(2_Part_1):209–220.

Martínez-Barajas, E., Delatte, T., Schluepmann, H., de Jong, G. J., Somsen, G. W., Nunes, C., Primavesi, L. F., Coello, P., Mitchell, R. A., and Paul, M. J. (2011). Wheat grain development is characterized by remarkable trehalose 6-phosphate accumulation pregrain filling: tissue distribution and relationship to snf1-related protein kinase1 activity. Plant physiology, 156(1):373–381.

Martinez-Carrasco, R. and Thorne, G. N. (1979). Physiological factors limiting grain size in wheat. Journal of experimental botany, 30(4):669–679.

McCouch, S. R., Wright, M. H., Tung, C.-W., Maron, L. G., McNally, K. L., Fitzgerald, M., Singh, N., DeClerck, G., Agosto-Perez, F., Korniliev, P., et al. (2016). Open access resources for genome-wide association mapping in rice. Nature communications, 7(1):1–14.

Meuwissen, T. H. E., Hayes, B. J., and Goddard, M. E. (2001). Prediction of total genetic value using genome-wide dense marker maps. Genetics, 157(4):1819–1829.

Morota, G. and Gianola, D. (2014). Kernel-based whole-genome prediction of complex traits: a review. Frontiers in Genetics, 5:363.

Obata, T. and Fernie, A. R. (2012). The use of metabolomics to dissect plant responses to abiotic stresses. Cellular and Molecular Life Sciences, 69(19):3225–3243.

Olsen, O.-A. (2004). Nuclear endosperm development in cereals and arabidopsis thaliana. The Plant Cell, 16(suppl_1):S214–S227.

Oszvald, M., Primavesi, L. F., Griffiths, C. A., Cohn, J., Basu, S. S., Nuccio, M. L., and Paul, M. J. (2018). Trehalose 6-phosphate regulates photosynthesis and assimilate partitioning in reproductive tissue. Plant Physiology, 176(4):2623–2638.

Peng, S., Huang, J., Sheehy, J. E., Laza, R. C., Visperas, R. M., Zhong, X., Centeno, G. S., Khush, G. S., and Cassman, K. G. (2004). Rice yields decline with higher night temperature from global warming. Proceedings of the National Academy of Sciences, 101(27):9971–9975.

Peng, S., Piao, S., Ciais, P., Myneni, R. B., Chen, A., Chevallier, F., Dolman, A. J., Janssens, I. A., Penuelas, J., Zhang, G., et al. (2013). Asymmetric effects of daytime and night-time warming on northern hemisphere vegetation. Nature, 501(7465):88–92.

Pérez, P. and de los Campos, G. (2014). Bglr: a statistical package for whole genome regression and prediction. Genetics, 198(2):483–495.

Pérez-Rodréguez, P. and de Los Campos, G. (2022). Multitrait bayesian shrinkage and variable selection models with the bglr-r package. Genetics, 222(1):iyac112.

Philipp, N., Weichert, H., Bohra, U., Weschke, W., Schulthess, A. W., and Weber, H. (2018). Grain number and grain yield distribution along the spike remain stable despite breeding for high yield in winter wheat. PLoS One, 13(10):e0205452.

Pielot, R., Kohl, S., Manz, B., Rutten, T., Weier, D., Tarkowská, D., Rolčík, J., Strnad, M., Volke, F., Weber, H., et al. (2015). Hormone-mediated growth dynamics of the barley pericarp as revealed by magnetic resonance imaging and transcript profiling. Journal of Experimental Botany, 66(21):6927–6943.

Ponnu, J., Wahl, V., and Schmid, M. (2011). Trehalose-6-phosphate: connecting plant metabolism and development. Frontiers in Plant Science, 2:70.

R Core Team (2022). R: A Language and Environment for Statistical Computing. R Foundation for Statistical Computing, Vienna, Austria.

Ren, R.-J., Wang, P., Wang, L.-N., Su, J.-P., Sun, L.-J., Sun, Y., Chen, D.-F., and Chen, X.-W. (2020). Os4bglu14, a monolignol β-glucosidase, negatively affects seed longevity by influencing primary metabolism in rice. Plant Molecular Biology, 104(4):513–527.

Riedelsheimer, C., Czedik-Eysenberg, A., Grieder, C., Lisec, J., Technow, F., Sulpice, R., Altmann, T., Stitt, M., Willmitzer, L., and Melchinger, A. E. (2012). Genomic and metabolic prediction of complex heterotic traits in hybrid maize. Nature genetics, 44(2):217–220.

Robin, X., Turck, N., Hainard, A., Tiberti, N., Lisacek, F., Sanchez, J.-C., and Möller, M. (2011). proc: an open-source package for r and s+ to analyze and compare roc curves. BMC bioinformatics, 12(1):1–8.

Schrag, T. A., Westhues, M., Schipprack, W., Seifert, F., Thiemann, A., Scholten, S., and Melchinger, A. E. (2018). Beyond genomic prediction: combining different types of omics data can improve prediction of hybrid performance in maize. Genetics, 208(4):1373–1385.

Smeekens, S. (2015). From leaf to kernel: trehalose-6-phosphate signaling moves carbon in the field. Plant Physiology, 169(2):912–913.

Sreenivasulu, N., Butardo Jr, V. M., Misra, G., Cuevas, R. P., Anacleto, R., and Kavi Kishor, P. B. (2015). Designing climate-resilient rice with ideal grain quality suited for high-temperature stress. Journal of Experimental Botany, 66(7):1737–1748.

VanRaden, P. (2008). Efficient methods to compute genomic predictions. Journal of Dairy Science, 91(11):4414–4423.

Vose, R. S., Easterling, D. R., and Gleason, B. (2005). Maximum and minimum temperature trends for the globe: An update through 2004. Geophysical Research Letters, 32(23).

Wada, H., Hatakeyama, Y., Onda, Y., Nonami, H., Nakashima, T., Erra-Balsells, R., Morita, S., Hiraoka, K., Tanaka, F., and Nakano, H. (2019). Multiple strategies for heat adaptation to prevent chalkiness in the rice endosperm. Journal of experimental botany, 70(4):1299–1311.

Wang, K., Li, Y., Wang, Y., and Yang, X. (2017). On the asymmetry of the urban daily air temperature cycle. Journal of Geophysical Research: Atmospheres, 122(11):5625–5635.

Wang, S., Wei, J., Li, R., Qu, H., Chater, J. M., Ma, R., Li, Y., Xie, W., and Jia, Z. (2019). Identification of optimal prediction models using multi-omic data for selecting hybrid rice. Heredity, 123(3):395–406.

Wase, N., Abshire, N., and Obata, T. (2022). High-throughput profiling of metabolic phenotypes using high-resolution gc-ms. In High-Throughput Plant Phenotyping, pages 235–260. Springer.

Welch, J. R., Vincent, J. R., Auffhammer, M., Moya, P. F., Dobermann, A., and Dawe, D. (2010). Rice yields in tropical/subtropical asia exhibit large but opposing sensitivities to minimum and maximum temperatures. Proceedings of the National Academy of Sciences, 107(33):14562–14567.

Westhues, M., Schrag, T. A., Heuer, C., Thaller, G., Utz, H. F., Schipprack, W., Thiemann, A., Seifert, F., Ehret, A., Schlereth, A., et al. (2017). Omics-based hybrid prediction in maize. Theoretical and applied genetics, 130(9):1927–1939.

Wheeler, T. and Von Braun, J. (2013). Climate change impacts on global food security. Science, 341(6145):508–513.

Xia, J., Chen, J., Piao, S., Ciais, P., Luo, Y., and Wan, S. (2014). Terrestrial carbon cycle affected by non-uniform climate warming. Nature Geoscience, 7(3):173–180.

Xing, Y., Zhang, Q., et al. (2010). Genetic and molecular bases of rice yield. Annual review of plant biology, 61(1):421–442.

Xu, S., Xu, Y., Gong, L., and Zhang, Q. (2016). Metabolomic prediction of yield in hybrid rice. The Plant Journal, 88(2):219–227.

Xu, Y., Xu, C., and Xu, S. (2017). Prediction and association mapping of agronomic traits in maize using multiple omic data. Heredity, 119(3):174–184.

Xu, Y., Zhao, Y., Wang, X., Ma, Y., Li, P., Yang, Z., Zhang, X., Xu, C., and Xu, S. (2021). Incorporation of parental phenotypic data into multi-omic models improves prediction of yield-related traits in hybrid rice. Plant biotechnology journal, 19(2):261–272.

Zhao, C., Liu, B., Piao, S., Wang, X., Lobell, D. B., Huang, Y., Huang, M., Yao, Y., Bassu, S., Ciais, P., et al. (2017). Temperature increase reduces global yields of major crops in four independent estimates. Proceedings of the National Academy of Sciences, 114(35):9326–9331.

Zhao, K., Tung, C.-W., Eizenga, G. C., Wright, M. H., Ali, M. L., Price, A. H., Norton, G. J., Islam, M. R., Reynolds, A., Mezey, J., et al. (2011). Genome-wide association mapping reveals a rich genetic architecture of complex traits in oryza sativa. Nature communications, 2(1):1–10.

Zhao, Y., Li, Z., Liu, G., Jiang, Y., Maurer, H. P., Würschum, T., Mock, H.-P., Matros, A., Ebmeyer, E., Schachschneider, R., et al. (2015). Genome-based establishment of a high-yielding heterotic pattern for hybrid wheat breeding. Proceedings of the National Academy of Sciences, 112(51):15624–15629.

Zhu, F., Paul, P., Hussain, W., Wallman, K., Dhatt, B. K., Sandhu, J., Irvin, L., Morota, G., Yu, H., and Walia, H. (2021). Seedextractor: an open-source gui for seed image analysis. Frontiers in plant science, 11:581546.

