## Supplemental File for "Evaluating metabolic and genomic data for predicting grain traits under high night temperature stress in rice"

### Tables

Table S1: List of metabolites

| Number | Metabolite | Number | Metabolite |
|---|---|---|---|
| 1 | hexanoic acid | 2 | alanine |
| 3 | valine | 4 | urea |
| 5 | ethanolamine | 6 | leucine |
| 7 | glycerol | 8 | nicotinic acid |
| 9 | isoleucine | 10 | proline |
| 11 | glycine | 12 | glyceric acid |
| 13 | citraconic acid | 14 | serine |
| 15 | threonine | 16 | Beta alanine |
| 17 | malic acid | 18 | dihydrouracil |
| 19 | threitol | 20 | methionine |
| 21 | aspartic acid | 22 | cytosine |
| 23 | trans-4-hydroxy proline | 24 | gamma-aminobutyric acid (GABA) |
| 25 | glutamic acid | 26 | hydroxybenzoic acid |
| 27 | asparagine | 28 | arabinose |
| 29 | lyxose | 30 | ribose |
| 31 | xylitol | 32 | arabitol |
| 33 | ribitol | 34 | diglycerol |
| 35 | 4-hydroxy-3-methoxybenzoic acid | 36 | glycerol-1 phosphate |
| 37 | glutamine | 38 | dihydroxybenzoic acid |
| 39 | ornithine | 40 | citrulline |
| 41 | citric acid | 42 | adenine |
| 43 | fructose-1 | 44 | allantoin-2 |
| 45 | altrose | 46 | lysine |
| 47 | histidine | 48 | glucose |
| 49 | tyrosine | 50 | mannitol |
| 51 | sorbitol | 52 | indoleacetic acid |
| 53 | pantothenic acid | 54 | glucosaminic acid |
| 55 | allantoin-3 | 56 | ferulic acid |
| 57 | N-acetyl-D-glucosamine | 58 | allo-inositol |
| 59 | tryptophan | 60 | 3.5-dimethoxy-4-hydroxycinnamic acid |
| 61 | uridine | 62 | eicosapentaenoic acid |
| 63 | adenosine | 64 | trehalose |
| 65 | maltose | 66 | sophorose |
| 67 | catechin | 68 | melibiose |
| 69 | isomaltose | 70 | galactinol |
| 71 | phosphoric acid | 72 | Sucrose |
| 73 | Raffinose | | |

### Figures

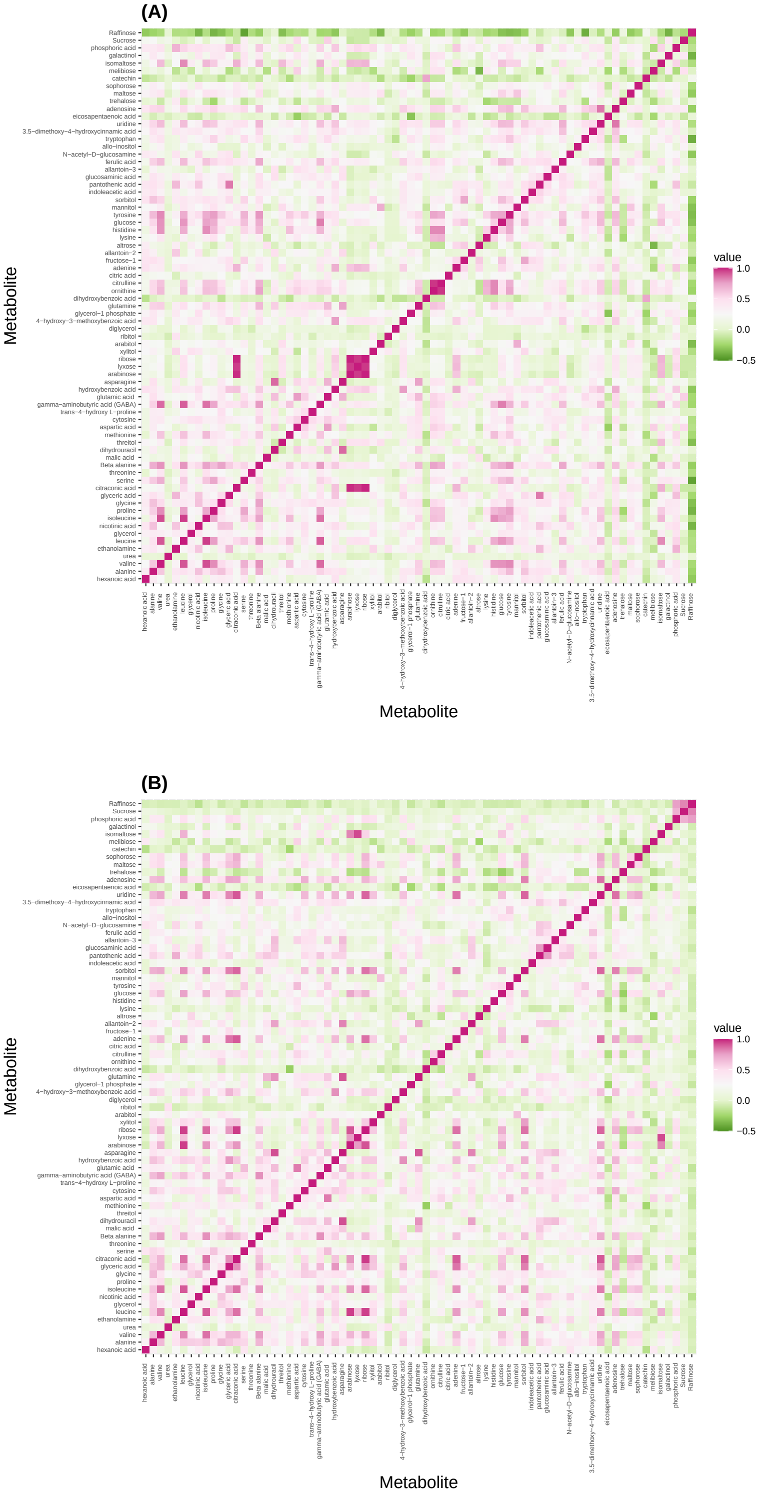


Figure S1: Pearson correlation heat map among metabolic profiles in control (A) and high night time temperature stress conditions (B).

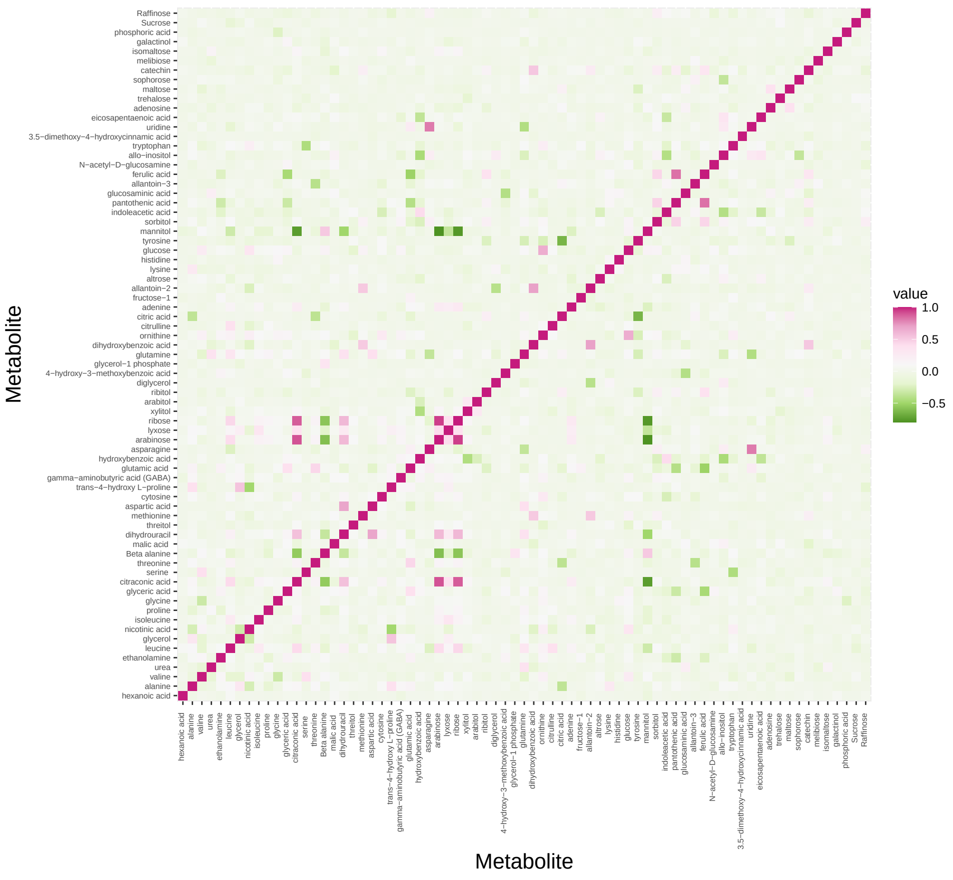


Figure S2: Pearson correlation heat map among metabolic profiles expressed as the ratio of control and high night time temperature stress.

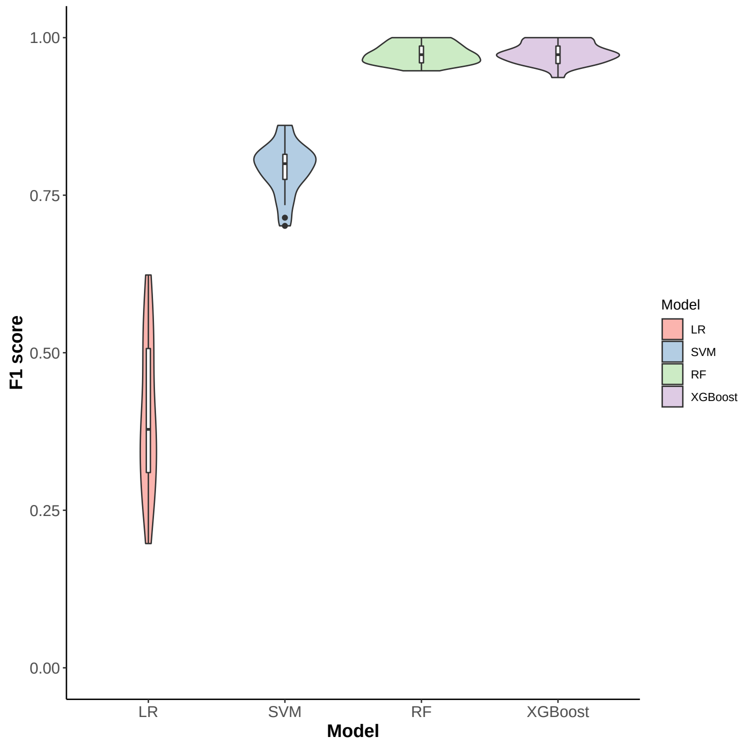


Figure S3: F1 scores of high night time temperature conditions (control and stress) using 73 metabolites. LR: logistic regression; SVM: support vector machine; and RF: random forest; XGBoost: extreme gradient boosting

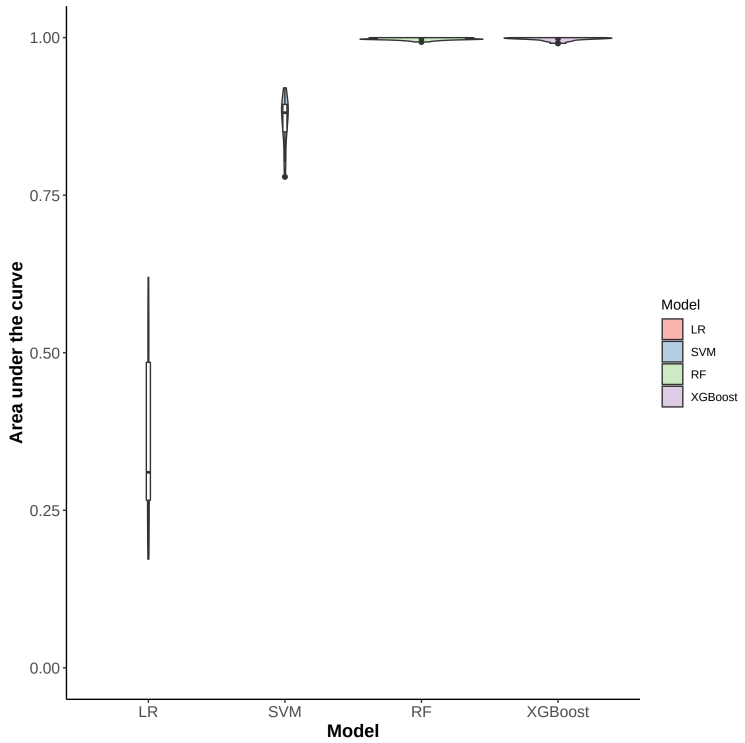


Figure S4: Area under the curve of high night time temperature conditions (control and stress) using 73 metabolites. LR: logistic regression; SVM: support vector machine; and RF: random forest; XGBoost: extreme gradient boosting

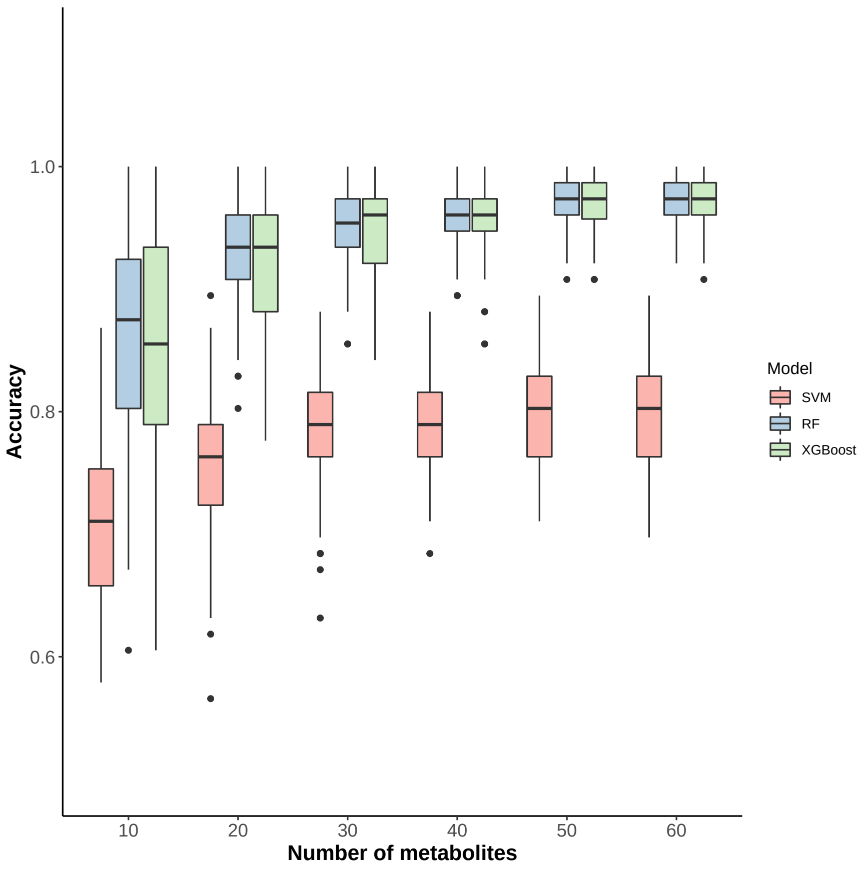


Figure S5: Classification accuracy of high night time temperature conditions (control and stress) using different number of metabolites using support vector machine (SVM), random forests (RF), and extreme gradient boosting (XGBoost).

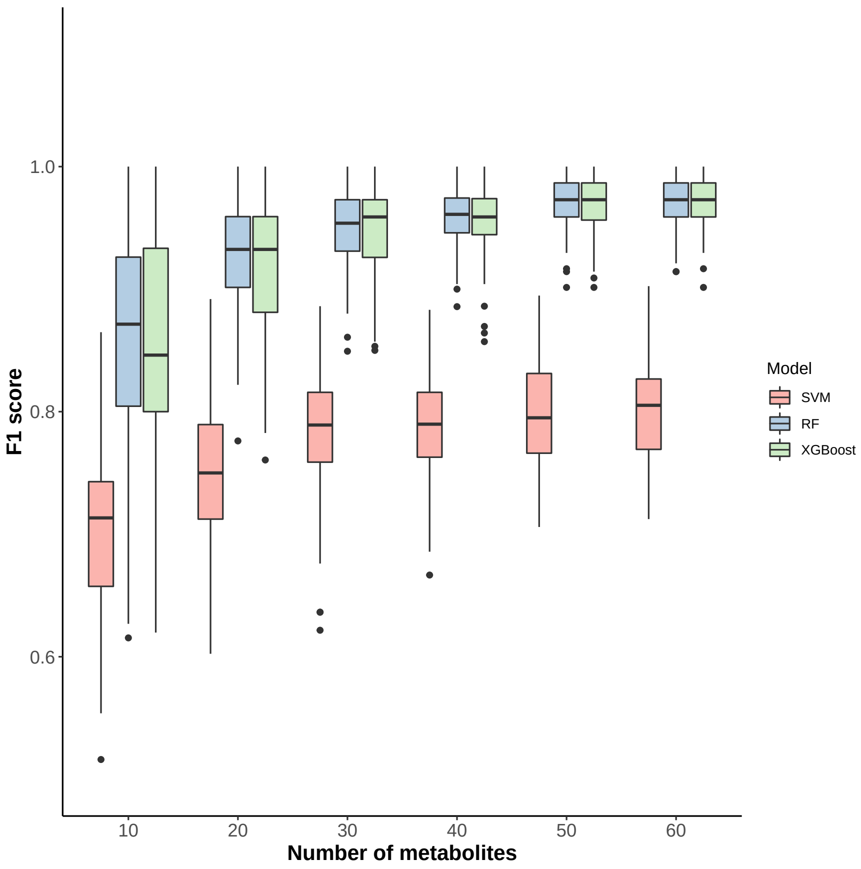


Figure S6: F1 scores of high night time temperature conditions (control and stress) using different number of metabolites using support vector machine (SVM), random forests (RF), and extreme gradient boosting (XGBoost).

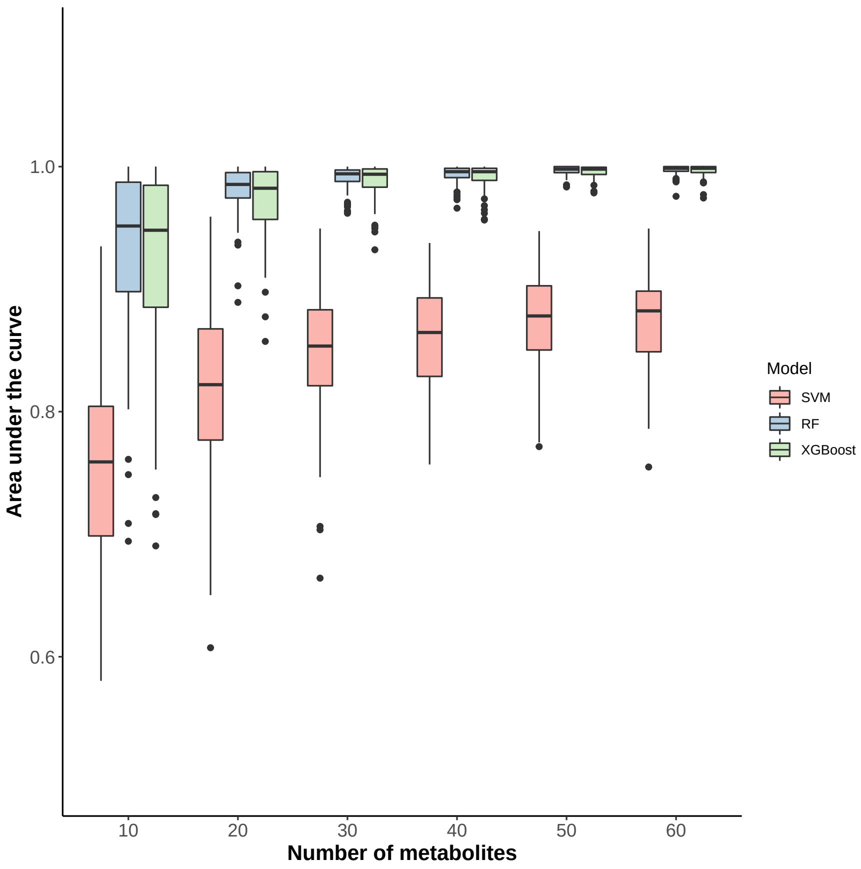


Figure S7: Area under the curve of high night time temperature conditions (control and stress) using different number of metabolites using support vector machine (SVM), random forests (RF), and extreme gradient boosting (XGBoost).

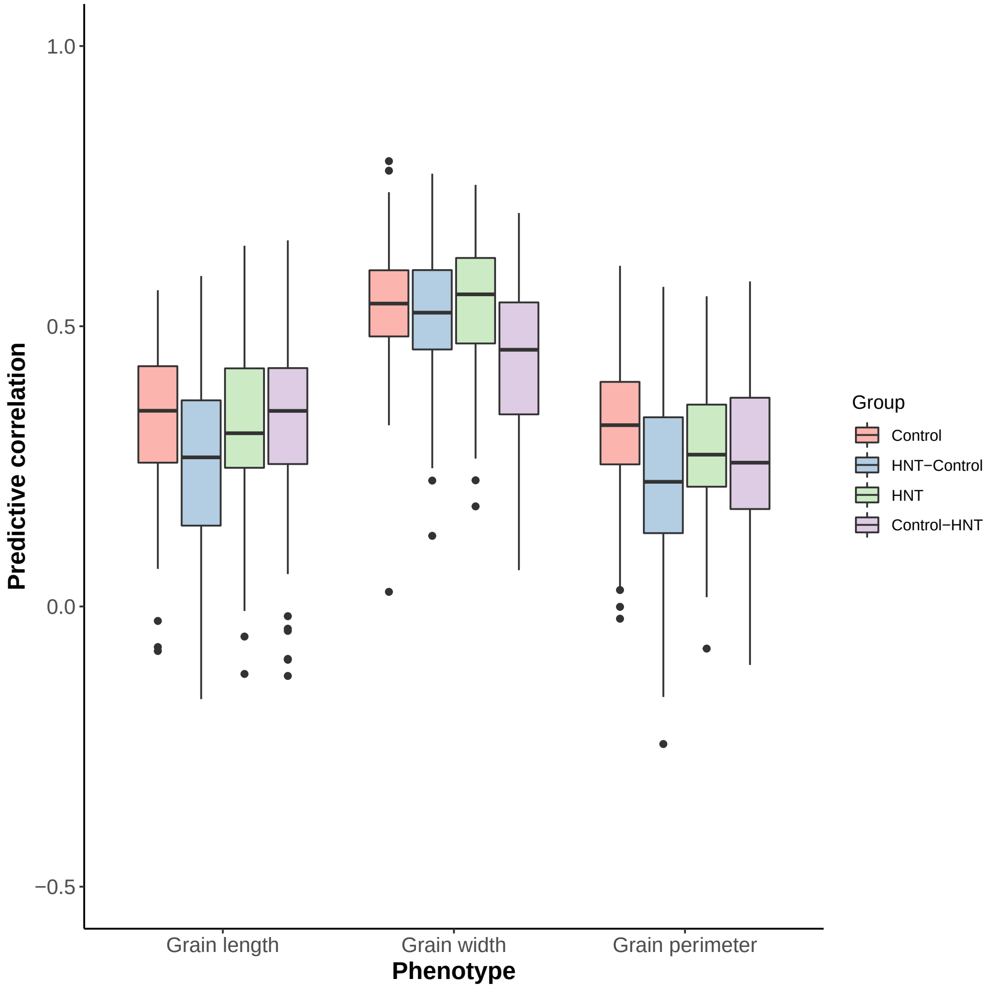


Figure S8: Metabolic predictions across high night time temperature (HNT) stress conditions. Control: Train the model using phenotypes and metabolites in control and predict control phenotypes from metabolites in control conditions. Control-HNT: Train the model using phenotypes and metabolites in control and predict HNT phenotypes from metabolites in HNT conditions using the metabolic effect estimated in control conditions. HNT: Train the model using phenotypes and metabolites in HNT and predict HNT phenotypes from metabolites in HNT conditions. HNT-Control: Train the model using phenotypes and metabolites in HNT and predict phenotypes from metabolites in control conditions using the metabolic effect estimated in HNT condition.
